# EGR4-dependent decrease of UTF1 is associated with failure to reserve spermatogonial stem cells in infertile men

**DOI:** 10.1101/2021.02.02.429371

**Authors:** Sara Di Persio, Tobias Tekath, Lara Marie Siebert-Kuss, Jann-Frederik Cremers, Joachim Wistuba, Xiaolin Li, Gerd Meyer zu Hörste, Hannes CA Drexler, Margot Julia Wyrwoll, Frank Tüttelmann, Martin Dugas, Sabine Kliesch, Stefan Schlatt, Sandra Laurentino, Nina Neuhaus

**Affiliations:** Centre of Reproductive Medicine and Andrology, University Hospital of Münster, Münster, 48149, Germany; Institute of Medical Informatics, University Hospital of Münster, Münster, 48149, Germany; Centre of Reproductive Medicine and Andrology, Department of Clinical and Surgical Andrology, University Hospital of Münster, Münster, 48149, Germany; Department of Neurology with Institute of Translational Neurology, University Hospital of Münster, Münster, 48149, Germany; Bioanalytical Mass Spectrometry Unit, Max Planck Institute for Molecular Biomedicine, Münster, 48149, Germany; Institute of Reproductive Genetics, University of Münster, Münster, 48149, Germany

**Keywords:** Male germline stem cells, spermatogonial stem cells, male infertility, impaired spermatogenesis, single cell RNA sequencing, human spermatogenesis, stem cell differentiation, testis.

## Abstract

Despite the high incidence of male infertility, about 70% of infertile men do not receive a causative diagnosis. To gain insights into the regulatory mechanisms governing human germ cell function in normal and impaired spermatogenesis (cryptozoospermic patients, crypto), we combined single cell RNA sequencing (>30.000 cells), proteome, and histomorphometric analyses of testicular tissues. We found major alterations in the crypto spermatogonial compartment with increased numbers of the most undifferentiated spermatogonia (PIWIL4^+^ State 0 cells). We also observed a transcriptional switch within the spermatogonial compartment driven by the increased and prolonged expression of the transcription factor *EGR4.* Intriguingly, EGR4-regulated genes included the chromatin-associated transcriptional repressor *UTF1*, which was downregulated. Histomorphometrical analyses showed that these transcriptional changes were mirrored at the protein level and accompanied by a change in the chromatin structure of spermatogonia. This resulted in a reduction of A_dark_ spermatogonia - characterized by tightly compacted chromatin and serving as reserve stem cells. These findings suggest that crypto patients are at a disadvantage especially in cases of gonadotoxic damage as they have less cells safeguarding the genetic integrity of the germline. We hypothesize that the more relaxed chromatin status of spermatogonia is dependent on decreased UTF1 expression caused by EGR4 activation. These identified regulators of the spermatogonial compartment will be highly interesting targets to uncover genetic causes of male infertility.

**One Sentence Summary:** Reserve spermatogonial stem cell depletion in infertile men is regulated by an EGR4-dependent UTF1 decrease, which changes chromatin morphology.

## Introduction

Infertility affects 10-15% of couples worldwide, with a male factor in half of the cases *(1*). Despite this high incidence of male infertility, about 70% of men do not receive a causative but only a descriptive diagnosis of impaired sperm production *(2*). This knowledge gap needs to be filled so that clinicians can counsel infertile men with regard to causal treatments, chances of sperm retrieval from testicular biopsies, transmission of infertility to the offspring, and potential health risks to the offspring and the males themselves. This latter aspect is of particular interest as male infertility and low sperm counts are associated with an increased risk of cancer, cardiometabolic disease and even premature mortality *(3*).

Of the infertile men, 7% are diagnosed with a severe form of oligoasthenoteratozoospermia termed “cryptozoospermia” (from the greek “kryptós”: hidden) as they have less than 0.1 million sperm in the ejaculate, rendering natural conception almost impossible *(2*). The surgical approach of testicular sperm extraction (TESE) often represents the only route to retrieve viable sperm for use in intracytoplasmic sperm injection (ICSI). Lower fertilization as well as implantation rates of transferred embryos *(4*) strongly indicate alterations not only in sperm production but also in the quality of the germ cells themselves. The underlying etiological factors for this severely impaired sperm production often remain unknown at cellular and molecular level but urgently need to be unveiled to improve clinical diagnostics, counselling, and prospectively treatment of infertile men.

Spermatogenesis is a highly complex process orchestrated by up to 10% of the male genome and regulated by a delicate interplay between the somatic environment and the germline. Spermatogenesis is supported by the spermatogonial stem cells (SSCs), a subpopulation of diploid spermatogonia, which undergo mitotic divisions, enter meiosis, and ultimately give rise to mature haploid sperm. SSCs are defined based on their functional properties, their ability to self-renew, and to give rise to differentiating germ cells. SSC systems in mice and men show species-specific differences *(5*). The human SSC system, according to the most accepted model, is classified as a progenitor-buffered system, containing quiescent reserve spermatogonia (A_dark_) in addition to self-renewing spermatogonia (A_pale_), the latter generating differentiating spermatogonia (B) *(6–8)*. These three different cell populations are classified based on their morphological appearance and the chromatin structure of their nuclei. In contrast to this, single cell RNA sequencing (scRNA-seq) analyses of the normal human testicular tissues enabled the identification of multiple spermatogonial states *(9–12)* indicative of a SSC compartment that is more heterogeneous than anticipated based on morphological properties. Furthermore, these studies indicated *PIWIL4* and *EGR4* as marker genes of the most undifferentiated spermatogonia *(9, 11)*. As the morphological appearance of these transcriptionally-defined spermatogonia remains largely unknown, it has not been possible so far to link the two classification systems. Importantly, the cellular and molecular alterations of the SSC compartment in stressful conditions such as male infertility, remain to be discovered. This will provide not only valuable information about human SSC biology but might also reveal the mechanisms behind the association of male infertility and cancer or a reduced life span *(3*). These insights will pave the way to understand the connection between male infertility and general health.

Furthermore, infertile men display alterations of the spermatogonial microenvironment, which has been suggested as a regulatory unit for spermatogonial fate decisions. Although the molecular and cellular properties of the human SSC-microenvironment remain largely unknown, an intact interplay between spermatogonia and their microenvironment must be considered a prerequisite for intact spermatogenesis and thereby fertility *(5*). Key players of the spermatogonial microenvironment include cells in direct physical contact with the spermatogonia (e.g. Sertoli and peritubular cells) and cells outside the seminiferous tubules (e.g. Leydig, endothelial, and perivascular cells and macrophages) *(5, 13)*.

In this study, we set a framework which combines scRNA-seq, proteomic, histomorphometric, and bioinformatical analyses to unveil the cellular and molecular changes in the spermatogonial compartment and its microenvironment of cryptozoospermic men. We revealed an increase in the number of PIWIL4^+^ undifferentiated spermatogonia and a depletion of the quiescent A_dark_ spermatogonia, considered as reserve stem cells. Moreover, we identified EGR4 as a regulator of these alterations. We found a proinflammatory microenvironment and identified FGF2-FGFR1/3 interaction as a communication channel between advanced germ cells and spermatogonia in cryptozoospermic testes. Intriguingly, our findings highlight alterations of the stem cell compartment and its regulators as an origin of impaired spermatogenesis, representing a milestone in the understanding of pathways underlying male infertility.

## Results

### Alterations of the spermatogonial compartment in cryptozoospermic men

To study cellular and molecular changes associated with male infertility, particularly in the spermatogonial compartment, we selected testicular biopsies from cryptozoospermic men (crypto) and normal controls (normal) based on clinical parameters (n=34, table S1). We used a multi-layered approach including quantitative histomorphometrical, scRNA-seq, and proteome analysis (Fig. 1A). Histological quantifications revealed a reduced proportion of tubules containing elongated spermatids as the most advanced germ cell type in the crypto group, while the percentage of tubules with round spermatids, spermatocytes, and spermatogonia remained unchanged (Fig. 1B). To assess the cellular composition of the spermatogonial compartment, we analyzed pan-spermatogonial marker MAGEA4 and undifferentiated spermatogonial marker UTF1 (Fig. 1C). We found a similar number of spermatogonia (MAGEA4^+^) per tubule in tissues from normal and crypto individuals (n=6 each). In contrast, tissues from the crypto group showed a reduction in UTF1^+^ spermatogonia (Fig. 1D and table S2).

**Fig. 1.**
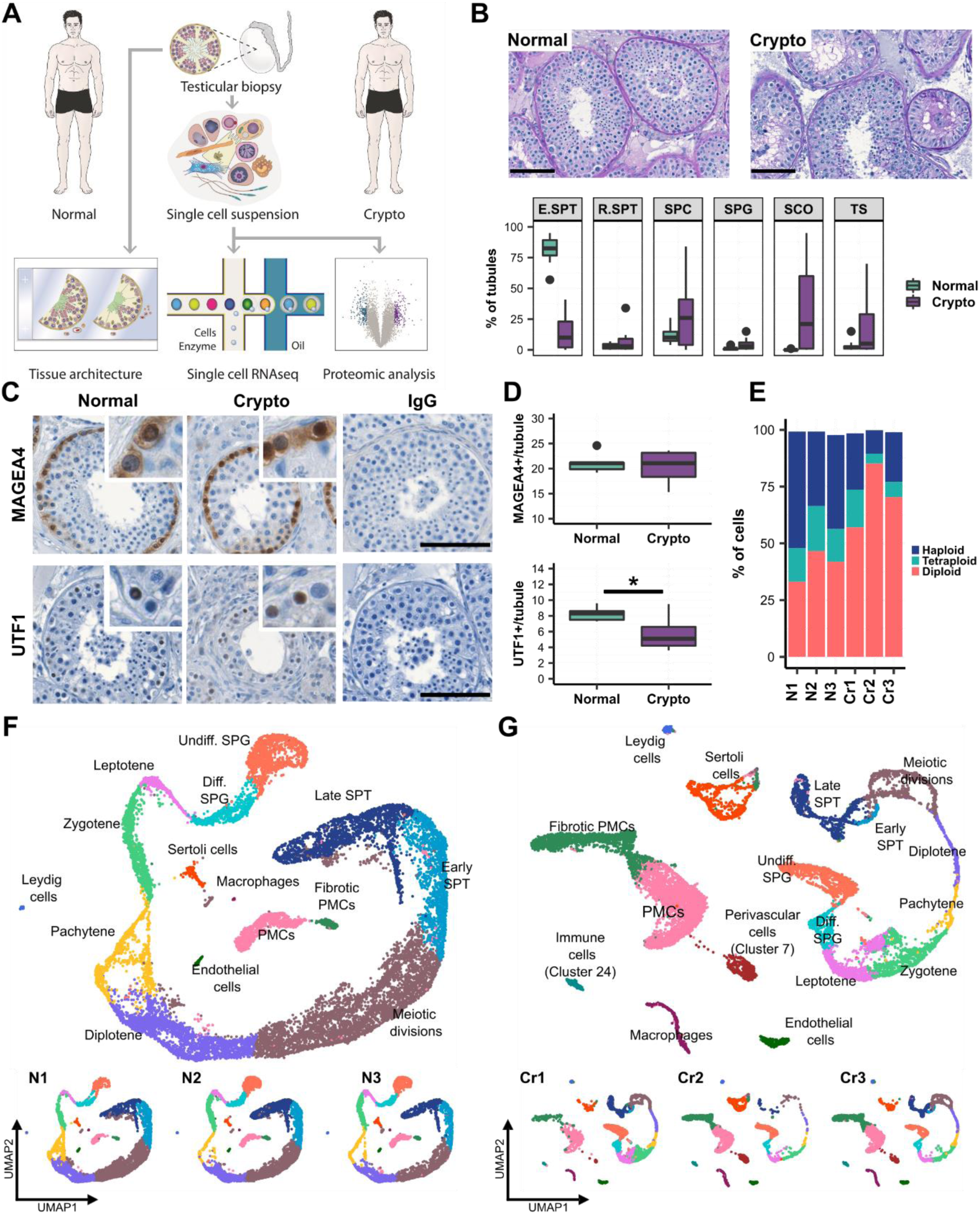
Crypto tissues show stable numbers of MAGEA4^+^ spermatogonia but reduced numbers of UTF1^+^ spermatogonia. (**A**) Experimental outline. (**B**) Upper panel: Representative Periodic acid-Schiff stained micrographs of the testicular tissue of one normal (left) and one crypto (right) sample. Scale bars = 100 µm. Lower panel: Box plots showing the percentages of tubules containing germ cells (most advanced germ cell type), only Sertoli cells, or tubular shadows in the patient cohort. Values per patient can be found in table S1. * p<0.05, *** p<0.001. (**C**) Micrographs showing the MAGEA4 and UTF1 staining in the normal and crypto testicular tissue. Inlays show examples of positive and negative cells for each staining. IgG negative controls show no staining. Scale bar = 100 μm. (**D**) Quantification of MAGEA4^+^ (upper) and UTF1^+^ (lower) spermatogonia per tubule in normal (n=6) and crypto samples (n=6). No statistical difference was found in the number of MAGEA4^+^ spermatogonia per tubule between normal and crypto samples. A significant reduction was found in the number of UTF1^+^ spermatogonia per tubule in the crypto samples (* p<0.05; Statistical details are available in table S2). (**E**) Stacked bar plots showing the ploidy of the single cell suspensions used for scRNA-seq analysis of each patient sample. (**F**) Upper panel: Uniform manifold approximation and projection (UMAP) plot of the integrated normal dataset. Clusters were assigned based on the expression of 55 markers genes (fig. S1B). Lower panel: scRNA-seq data from the 3 normal samples. Each color represents a different cell type. (**G**) Upper panel: UMAP plot of the integrated crypto dataset. The clusters were assigned by anchoring integration using the normal dataset as reference. Lower panel: scRNA-seq data from the 3 crypto samples.

In-depth characterization of the molecular changes was performed by scRNA-seq analysis of normal and crypto testicular biopsies (n=3 each). Suitable samples were selected based on ploidy analysis, ensuring presence of haploid spermatids in all samples (Fig. 1E). To exclude genetic causes for male infertility we performed whole exome sequencing analysis. No relevant variants were identified in any of the three cryptozoospermic men rendering the potential genetic cause for their infertility unidentifiable for the time being and making a uniform cause highly unlikely. Following scRNA-seq and quality control, data from 15 546 and 13 144 cells were ultimately included in the analysis for normal and crypto samples, respectively (table S3). Unsupervised clustering resulted in a comparable number of clusters (30 versus 29) between the two groups (fig. S1, A and B). To assign identities to the clusters while still considering eventual biological differences between the groups, we used a two-step approach. Firstly, we used 55 published marker genes to identify 15 cell types in the normal dataset (fig. S1B) *(9– 12)*. Notably, the latent space replicated the known differentiation process from spermatogonia to late spermatids (spiral-like shape in Fig. 1F). Using this as reference, we projected the same labels *(14*) onto the crypto dataset and found the expected cell types represented (Fig. 1G). Correct cluster assignment was confirmed by evaluating the expression of specific marker genes in the crypto dataset (fig. S1C). Importantly, all samples contributed to all cell type clusters in each dataset (Fig. 1, F and G). Additionally, using marker gene expression, we identified two additional clusters in the crypto group without equivalence in the normal dataset namely, perivascular cells (cluster 7) and immune cells (cluster 24; Fig. 1G).

### Cellular and transcriptional changes in germ and somatic cell compartments

To explore the similarities and differences between the two groups, we combined all normal and crypto datasets (Fig. 2A). Consistent with the histomorphometrical analysis (Fig. 1D) evaluation of the scRNA-seq dataset showed comparable percentages of spermatogonia in both groups (Fig. 2B). In fact, percentages were similar from spermatogonia up to zygotene spermatocytes (Fig. 2B). However, from the pachytene spermatocyte stage onwards, the crypto group displayed a striking reduction in germ cells (Fig. 2B). This was corroborated by parallel quantitative proteomic analysis, showing comparable levels of spermatogonial marker MAGEA4 and reduced expression of markers for leptotene/pachytene spermatocytes (DPH7, PIWIL1) as well as spermatids (PRM2; Fig. 2C). To assess the changes at the transcriptional level between all the identified cell types, we performed differential gene expression (DGE) analysis (table S4) and identified genes uniquely differentially expressed in each germ cell stage (fig. S2 and table S5). Interestingly, *UTF1* and *EGR4* were down- and up-regulated respectively in the crypto undifferentiated spermatogonia (fig. S2A). Curiously, genes associated with the gene ontology (GO) terms “Cell cycle”, “Chromosome organization”, “DNA repair”, and “Telomere organization” (e.g. *BRCA2*, *CENPA*, *SMC1A*, *SYCP3*, *MEIOB*, *CETN2*) were differentially expressed in differentiating spermatogonia compared to pachytene spermatocytes (fig. S2, B to E).

**Fig. 2.**
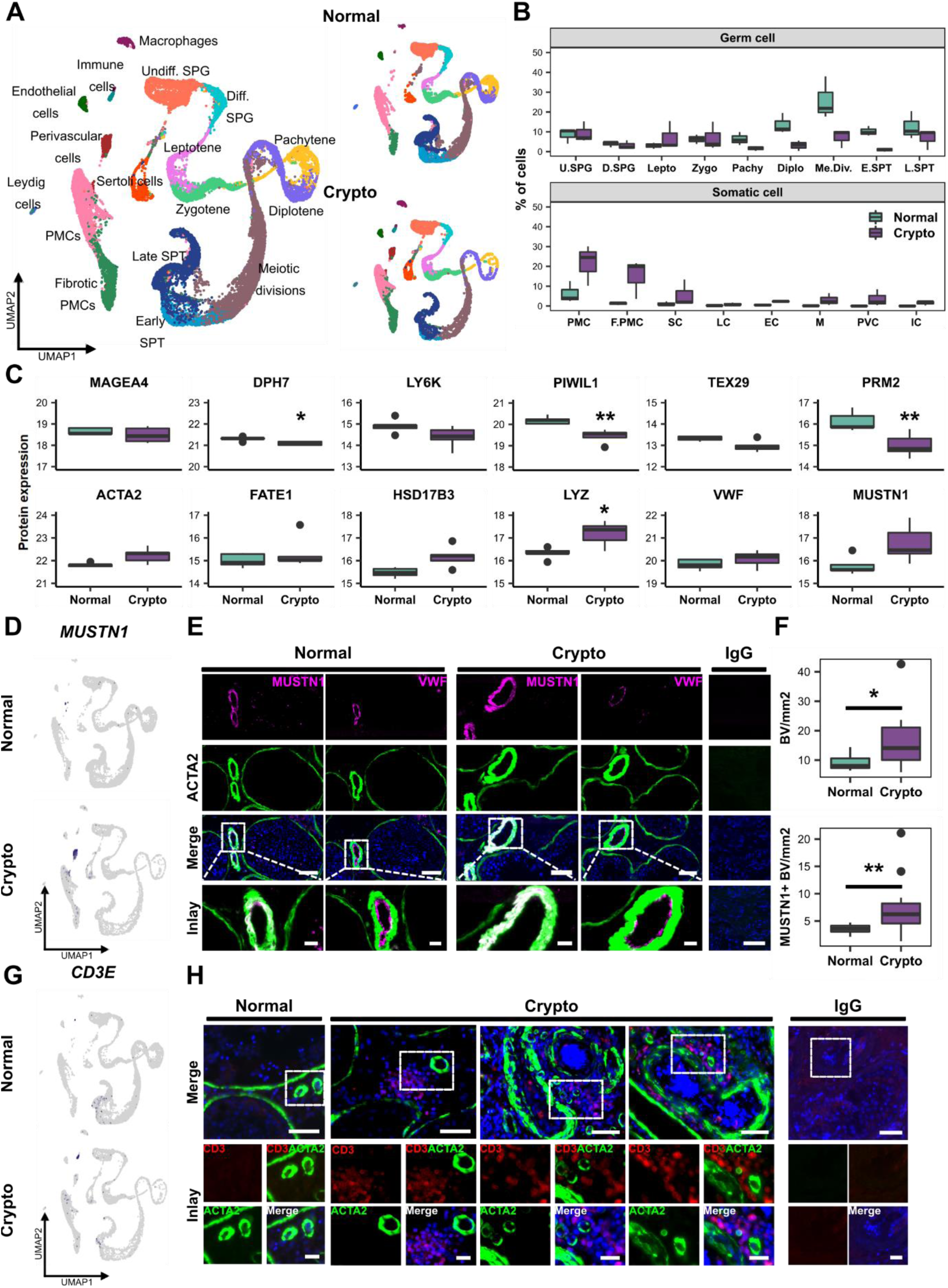
Exploration of cellular and transcriptional changes in crypto testicular tissues. **(A)** Left panel: UMAP plot of the all integrated dataset. Right panel: Contribution of the normal (15 546 cells) and crypto (13 144 cells) datasets to the all integrated dataset. The cells are color coded according to the respective cell types. **(B)** Box plot comparing the germ and somatic cell proportions in normal and crypto scRNA-seq datasets. **(C)** Panel of box plots representing the protein expression of 12 marker genes used in the cluster assignment (MAGEA4: Spermatogonia; DPH7: Leptotene spermatocytes; LY6K: Zygotene spermatocytes; PIWIL1: Pachytene spermatocytes; TEX29: Early spermatids; PRM2: Elongated spermatids; ACTA2: Peritubular myoid cells (PMCs); FATE1: Sertoli cells; HSD17B3: Leydig cells; VWF: Endothelial cells; LYZ: Macrophages; MUSTN1: Perivascular cells). Protein expression was measured performing bottom-up mass spectrometry analysis on single cell suspensions (normal n=5, crypto n=4 patients) and shows a reduction of germ cells, starting from the leptotene spermatocytes, and an increase of macrophages in crypto patients. * p<0.05, ** p<0.01. **(D)** Feature plots highlighting the expression of *MUSTN1* (Perivascular cell marker gene) in normal and crypto datasets. **(E)** Representative micrographs showing blood vessels in consecutive sections of normal and crypto testicular tissues stained for MUSTN1 (magenta)/ACTA2 (green) and for VWF (magenta)/ACTA2 (green). The tissue sections were counterstained with DAPI (blue). Double positive MUSTN1/ACTA2 cells were found surrounding the endothelial layer of the blood vessels. The IgG control showed no immunological staining. Scale bar = 100 μm (main) and 20 μm (inlays). **(F)** Box plots representing the number of ACTA2^+^ blood vessels (Upper) and MUSTN1^+^ blood vessels (lower) per mm^2^ of tissue in the normal (n=12) and crypto samples (n=13). A significant increase of blood vessels and MUSTN1^+^ blood vessels was found between the two cohorts (Statistical details are available in table S2). **(G)** Feature plots highlighting the expression of *CD3E* (Immune cell marker gene) in normal and crypto datasets. **(H)** Representative micrographs showing CD3^+^ immune cells (Red) and ACTA2^+^ blood vessels (Green) in normal and crypto testicular tissues. The tissue sections were counterstained with DAPI (Blue). CD3^+^ immune cells were found solely in the crypto group in close proximity to blood vessels. The IgG control showed no immunological staining. Scale bar = 50 μm (main) and 20 μm (inlays).

In the somatic cell compartment, the most striking difference was the increase in the proportions of peritubular myoid cells (PMCs), fibrotic PMCs, and macrophages (Fig. 2B); the latter finding in particular was supported by proteomic data, which revealed significantly increased macrophage marker LYZ expression (Fig. 2C). Among somatic cells, PMCs showed the most profound changes in DGE analysis, with 83 down- and 149 up-regulated genes, respectively, in the crypto group. GO analysis revealed an enrichment of genes involved in “extracellular matrix components and organization” (fig. S3A and table S5). Because perivascular and immune cells did not have equivalents in the normal dataset, no direct comparison was conducted between normal and crypto samples (fig. S3B). DGE analysis between perivascular cells and PMCs in the crypto dataset revealed 48 genes (fig. S3C and table S6) regulating muscle contraction (fig. S3D). To localize perivascular cells at tissue level we selected the musculoskeletal marker *MUSTN1* from the upregulated genes, which was exclusively expressed in this cluster at RNA level (Fig. 2D). MUSTN1 was specifically expressed in blood vessels, where it co-localized with ACTA2. Indeed, these ACTA2^+^/MUSTN1^+^ cells surrounded VWF^+^ endothelial cells (Fig. 2E). Quantification of the total number of blood vessels/mm^2^ revealed a significantly higher number of blood vessels in crypto samples (Fig. 2F and table S2). Similar results were obtained for the number of MUSTN1^+^ blood vessels/mm^2^ (Fig. 2F and table S2). We therefore concluded that MUSTN1 is a specific perivascular cell marker and that an increased proportion of MUSTN1^+^ blood vessels is a specific feature of crypto samples. DGE analysis comparing immune cells to all other crypto cells resulted in 195 differentially expressed genes, including T-cell marker *CD3E* (Fig. 2G, fig. S3E and table S6). To evaluate which types of immune cells were present in crypto testes, we performed pathway analysis and found involvement of cytotoxic T lymphocyte pathways (fig. S3F). Immunofluorescence analysis localized CD3^+^ T cells in the vicinity of blood vessels in crypto samples but not in the normal (Fig. 2H). To investigate the crosstalk between spermatogonia and their microenvironment, we used CellphoneDB *(15*). We found 50 significant ligand-receptor interactions considering both datasets (Fig. 3). A significant interaction was detected between the spermatogonial-based receptors *FGFR1* and *3* and the ligand *FGF2* produced by pachytene and diplotene spermatocytes in the crypto group. Moreover, spermatogonial-located *ACKR2* showed significant interaction with its ligands secreted by endothelial cells, perivascular cells, macrophages, and immune cells (*CCL2*, *CCL3*, *CCL4*, *CCL5, CCL3L1* and *CCL14*) in the crypto datasets but not in the normal.

**Fig. 3.**
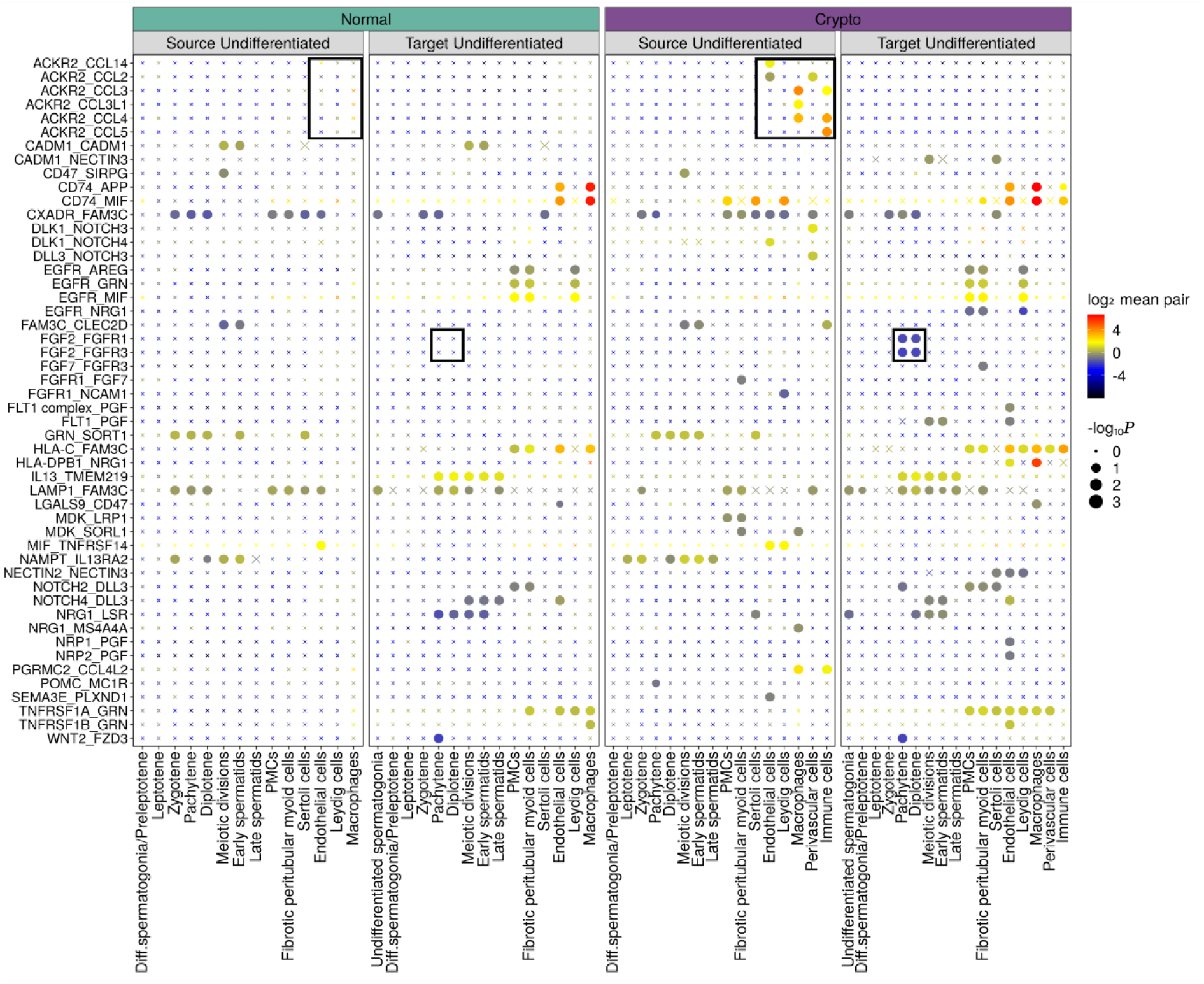
Ligand-receptor interaction analysis. Summary of ligand–receptor interactions between the normal and crypto undifferentiated spermatogonia and the other cell types in the tissues. P–values are represented by the size of each circle. The color gradient indicates the level of interaction. Black squares indicate selected interactions that show a different significance in the normal and crypto datasets.

### Crypto undifferentiated spermatogonia activate the EGR4 regulatory network

SCENIC was used to identify the gene regulatory network changes in the crypto somatic and germ cell populations *(16*). This pipeline identifies groups of genes, named regulons, co-regulated by a transcription factor. In total, in the two datasets we identified 403 regulons, each containing up to 2 450 genes (table S7). To identify differentially activated regulons between the normal and the crypto dataset, we selected the 15 regulons with the highest specificity score for each cluster in the normal dataset. We then focused on those regulons in the crypto dataset that showed at least 20% change in the proportion of cells per cluster in which that specific regulon was active (Fig. 4A and table S8). Comparative evaluation of the binarized AUC (area under the curve) score of each regulon revealed 13 regulatory networks that were differentially activated between the two datasets (bold in Fig. 4A). As perivascular and immune cells were uniquely represented in the crypto dataset, they were not included in this analysis. In the somatic compartment, although PMC and fibrotic PMC clusters presented the most striking differences in cell proportions, no regulon was differentially activated in these cells. Similarly, we could not find differentially activated regulons for Leydig cells. However, one differentially activated regulon was found in the macrophages in the crypto dataset (activation of RUNX3, 75 genes, fig. S4).

**Fig. 4.**
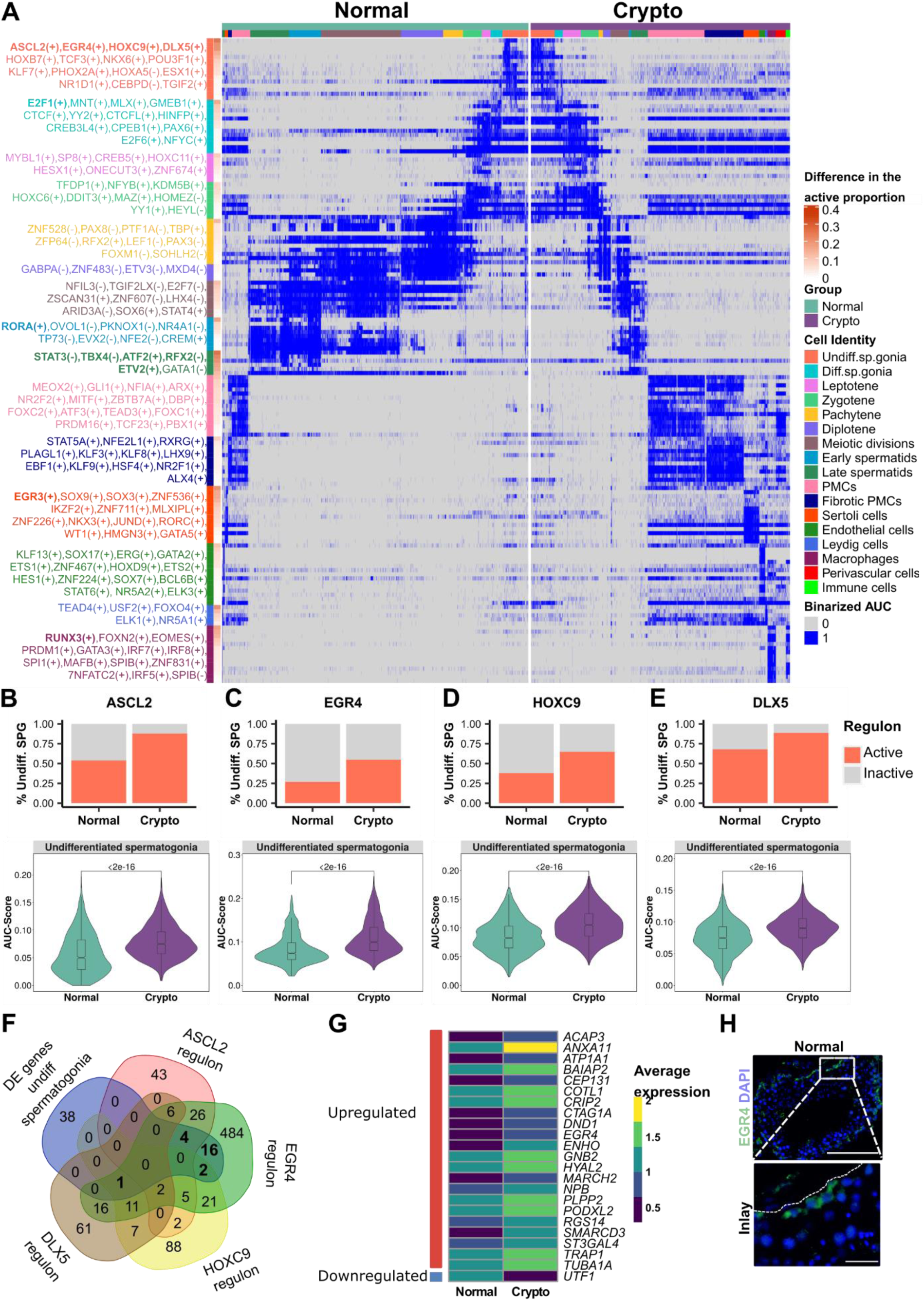
Undifferentiated spermatogonia activate EGR4 regulatory network in the crypto group. (**A**) Binarized double heatmap showing the AUC score (area under the recovery curve; it scores the activity of regulons) of the identified regulons plotted separately for the normal (teal, left) and crypto (purple, right) datasets. For each cellular cluster, the first 15 regulons showing the highest regulon specificity score (RSS) in the normal dataset were plotted. The regulons written in bold are those showing a difference of at least 20% in the proportion of cells with the active regulon in the crypto dataset and a significant difference in the regulon specific AUC score between normal and crypto datasets. (**B-E**) Upper panel: Stacked bar plots comparing the proportion of undifferentiated spermatogonia with active (**B**) ASCL2, (**C**) EGR4, (**D**) HOXC9 and (**E**) DLX5 regulons. Lower panel: Violin plots comparing the AUC score of the (**B**) ASCL2, (**C**) EGR4, (**D**) HOXC9 and (**E**) DLX5 regulons in the normal and crypto undifferentiated spermatogonia. A significant increase was found for all four regulons in the crypto dataset (BH corrected p-value of Mann-Whitney U test). A complete list of the regulons and their RSS and AUC scores is available in table S8. (**F**) Overlap among the 61 genes uniquely differentially expressed in undifferentiated spermatogonia and the genes putatively regulated by ASCL2, EGR4, HOXC9 and DLX5. (**G**) Heatmap showing the expression in the normal and crypto undifferentiated spermatogonia of the 23 differentially expressed genes putatively regulated by EGR4. (**H**) Micrographs showing the expression of EGR4 in spermatogonia in normal testicular tissue. Scale bars: main=100 µm; inlay=20 µm.

Contrary to these minor changes in somatic cells, germ cells showed deeper alterations in their gene regulatory networks. Undifferentiated spermatogonia displayed four regulons with significantly enhanced activation in the crypto samples: ASCL2 (88 genes), EGR4 (594 genes), HOXC9 (139 genes), and DLX5 (104 genes) (Fig. 4B-E). Intriguingly, among the genes regulated by EGR4, we identified *ASCL2*, *HOXC9*, and *DLX5* whereas none of the other transcription factors seemed to regulate the expression of *EGR4*. Comparison between the regulon and the DGE analysis of the undifferentiated spermatogonia showed that 23 of the differentially expressed genes were predicted to be regulated by EGR4 (Fig. 4, F and G). Follow-up histological analysis showed specific expression of EGR4 in spermatogonia (Fig. 4H). Moreover, the E2F1 regulon (685 genes) showed a significant activation in differentiating spermatogonia (fig. S4A). While no regulons in the crypto dataset were differentially active specifically in spermatocytes, early spermatids showed activation of RORA (fig. S4B), and late spermatids showed deactivation of STAT3, TBX4, ATF2, and RFX2, and activation of ETV2 (fig. S4, C to G).

### Crypto spermatogonial compartment shows an increased number of PIWIL4^+^ spermatogonia

To further scrutinize the changes in the spermatogonial compartment, we identified specific spermatogonial subtypes based on marker genes reported in existing single cell studies *(9, 11)*. For this, we subset and re-clustered undifferentiated and differentiating spermatogonial clusters from both datasets (fig. S5A). Upon analyzing the expression of the different published markers in the normal dataset, we could assign the clusters and identify six spermatogonial states: State 0 (*PIWIL4, PHDGH, EGR4*), State 0A (*UTF1, SERPINE2, FGFR3*), State 0B (*NANOS2*), State 1 (*GFRA1, GFRA2, NANOS3, ID4*), State 2 (*KIT, MKI67, DMRT1, DNMT1*), and State 3 (*STRA, SYCP3*) (fig. S5B and Fig. 5A). Notably, the markers mentioned display the highest, but not unique, expression levels in the respective states. Using the data from normal spermatogenesis as reference, these labels were projected onto the crypto dataset (Fig. 5B). For further analyses, both datasets were integrated (Fig. 5C). To compare our assignment with those published by Guo et al., and Sohni et al., we performed correlation analysis among the three datasets *(9, 11)*. States 0, 0A and 0B in our assignment showed the highest correlation with State 0 in Guo et al. (fig. S5C) and with SSC1-B,-A, and -C respectively in Sohni et al. (fig. S5D).

**Fig. 5.**
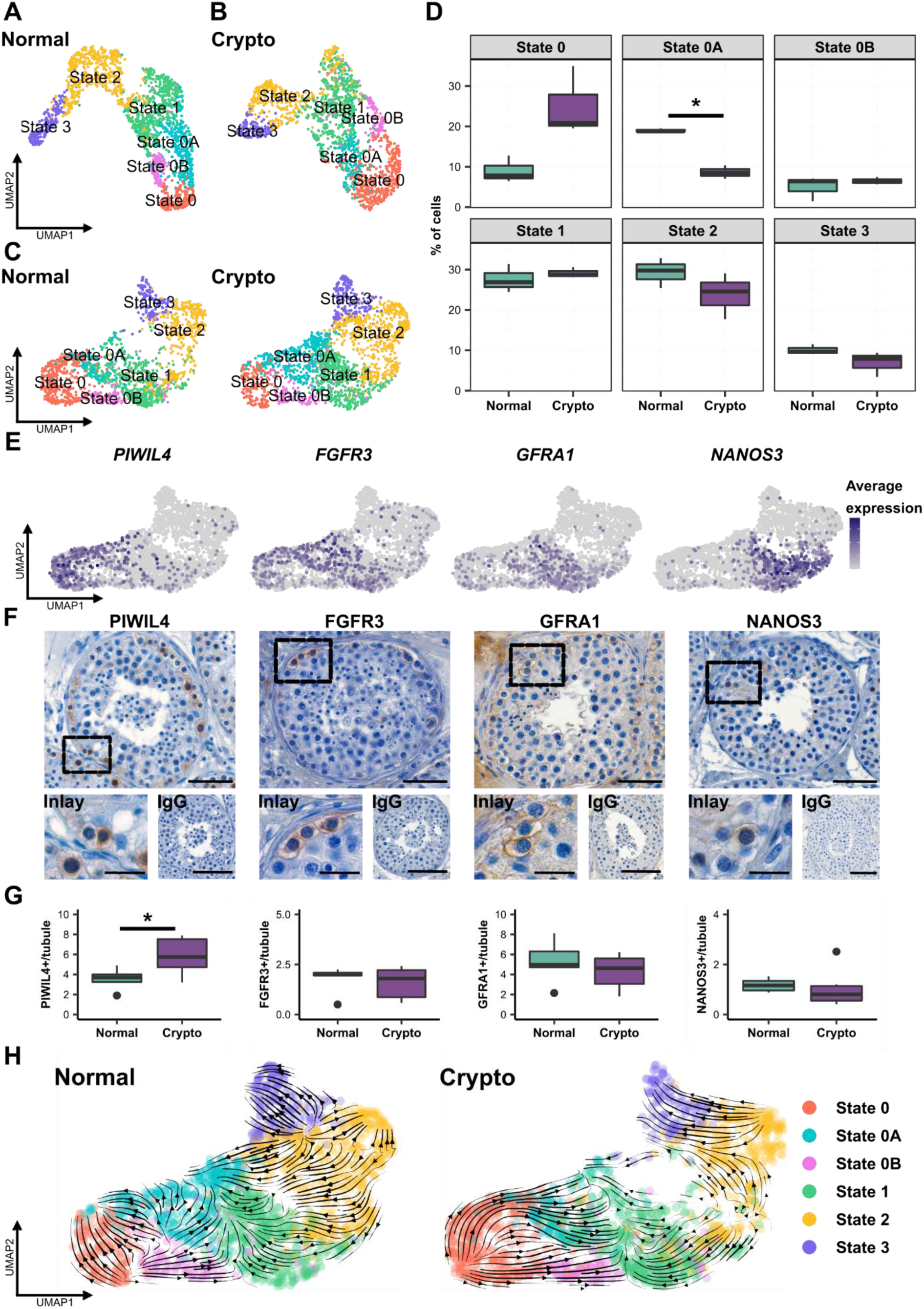
The crypto SSC compartment shows altered composition with an increased number of PIWIL4^+^ spermatogonia. (**A**) UMAP plot showing the cluster assignment of the normal dataset. Six clusters were assigned based on the expression of markers shown in fig. S5B: State 0, 0A, 0B, 1, 2, and 3. Each color represents a different cluster. (**B**) UMAP plot showing the cluster assignment of the crypto dataset. The clusters were assigned by the anchoring integration method using the normal dataset as reference. Each color represents a different cluster, as in the normal dataset. (**C**) UMAP plot showing the integrated normal and crypto spermatogonial datasets. Each color represents a different cluster, as in the normal dataset. (**D**) Box plots representing the different cell proportions in the normal and crypto spermatogonial datasets. The proportion of State 0A spermatogonia was significantly reduced in the crypto dataset (* p<0.05). (**E**) Feature plots showing the expression level of *PIWIL4, FGFR3, GFRA1*, and *NANOS3* in the all integrated datasets. (**F**) Micrographs showing the PIWIL4, FGFR3, GFRA1 and NANOS3 staining in the testicular tissue. Inlays show examples of positive and negative cells for each staining. IgG negative controls show no staining. Scale bars: main = 50 μm, Inlay=20 μm; IgG=100 μm. (**G**) Quantification of PIWIL4^+^, FGFR3^+^, GFRA1^+^, and NANOS3^+^ spermatogonia per tubule in normal (n=6) and crypto samples (n=6). A significant increase was found in the number of PIWIL4^+^ spermatogonia per tubule in the crypto samples (* p<0.05). No statistical difference was found in the total number of GFRA1^+^, FGFR3^+^ and NANOS3^+^ spermatogonia between normal and crypto samples. Statistical details are available in table S2. (**H**) RNA velocities derived from the scVelo dynamical model for the normal and crypto spermatogonial dataset are visualized as streamlines in UMAP plots. A change in the arrow directionality can be observed in the crypto dataset.

We assessed the cellular composition of the spermatogonial compartment and found an increased number of cells in State 0 and a significantly decreased proportion of cells in State 0A in the crypto dataset. There were no differences in cell numbers in any other states (Fig. 5D).

To reveal whether this shift in spermatogonial states was also represented at the cellular level, and considering that the number of UTF1^+^ spermatogonia is significantly decreased (Fig. 1, C and D), we analyzed *PIWIL4*, *FGFR3*, *GFRA1,* and *NANOS3* (Fig. 5, E and F) using quantitative histomorphometric analysis. The altered proportions of the spermatogonial states identified based on scRNA-seq data were largely corroborated by cell quantification. We found a significant increase in PIWIL4^+^ spermatogonia (State 0) per seminiferous tubule, and no significant changes in FGFR3^+^, GFRA1^+^, and NANOS3^+^ spermatogonia (Fig. 5G and table S2). To further unravel the alterations of the spermatogonial compartment, RNA velocity analysis *(17*) was performed setting State 0 as the origin of the spermatogenic differentiation process. *(9*) The velocity streamlines (which depend on the mRNA maturation in each cell) indicate that States 0 and 0A in the normal samples are transcriptionally not directed toward State 1, unlike State 0B (Fig. 4I). In contrast, in the crypto samples the velocity streamlines in States 0, 0A, and 0B appear to be preferentially oriented toward State 1 (Fig. 5H). This was evident in all three crypto samples (fig. S5E).

### The crypto spermatogonial compartment shows reduced numbers of A_dark_ spermatogonia

To evaluate the expression profile of the morphologically-defined A_dark_ spermatogonia, we assessed how many A_dark_ spermatogonia were PIWIL4^+^, UTF1^+^, FGFR3^+^, GFRA1^+^, or NANOS3^+^. Semi-quantitative evaluation in tissues showed the majority of A_dark_ spermatogonia in the control group were UTF1^+^ (78.4%), whereas only 5.8% were PIWIL4^+^, 5.7% were FGFR3^+^, 18.1% were GFRA1^+^, and no A_dark_ cells were NANOS3^+^ (Fig. 6, A to D).

**Fig. 6.**
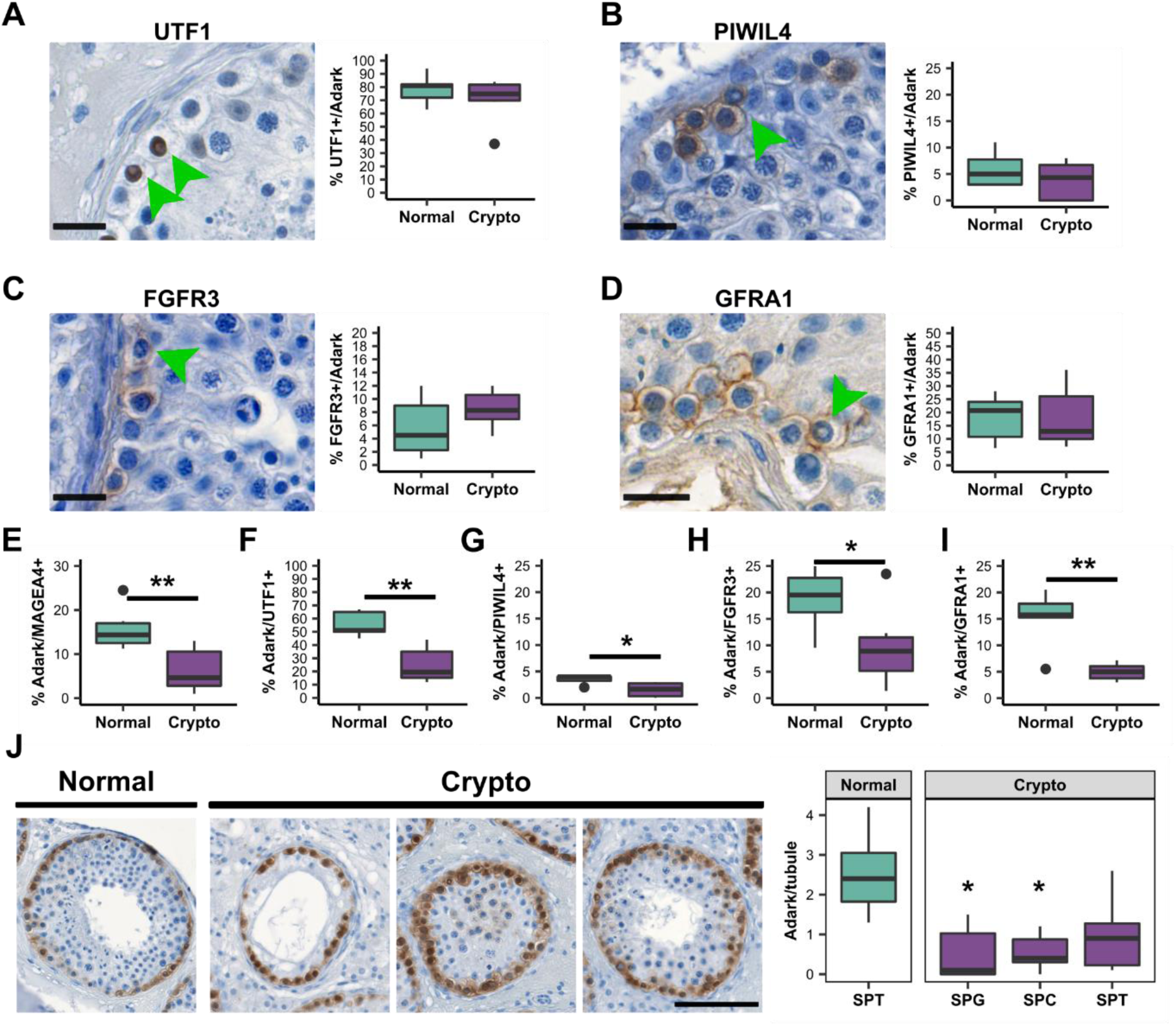
The crypto group shows altered numbers of A_dark_ spermatogonia. (**A-D**) Right panel: Micrographs showing (**A**) UTF1^+^, (**B**) PIWIL4^+^, (**C**) FGFR3^+^, (**D**) GFRA1^+^ A_dark_ spermatogonia. Green arrow heads indicate A_dark_ spermatogonia. Scale bars=20 µm. Left panel: Boxplots showing the percentage of UTF1^+^ (**A**), PIWIL4^+^ (**B**), FGFR3^+^ (**C**) and GFRA1^+^ (**D**) spermatogonia among the A_dark_ population in normal (n=6) and crypto samples (n=6). A mean of 78.4% of the A_dark_ spermatogonia were UTF1^+^, 5.8% were PIWIL4^+^, 5.7% were FGFR3^+^ and 18.1% were GFRA1^+^ in the normal dataset. No significant difference was found in the crypto samples. (**E-I**) Quantification of the percentage of A_dark_ spermatogonia among MAGEA4^+^ (**E**), UTF1^+^ (**F**), PIWIL4^+^ (**G**), FGFR3^+^ (**H**) and GFRA1^+^ (**I**) spermatogonial populations in normal (n=6) and crypto samples (n=6). A significant reduction of A_dark_ spermatogonia was found among all the cell types (**p <0.01, *p <0.05). (**J**) Quantification of the proportion of A_dark_ spermatogonia per tubule according to the most advanced germ cell type present. Left: Representative micrographs showing in the order one tubule from a normal sample containing spermatids as the most differentiated germ cell and three tubules from a crypto sample containing either spermatogonia (SPG), spermatocytes (SPC), or spermatids (SPT) as the most differentiated germ cell. Scale bar = 100µm. Right panel: Boxplot showing the quantification of A_dark_ spermatogonia per tubule in normal (n=6) and crypto samples (n=6). *p <0.05. The asterisks refer to the comparison with the normal samples. Statistical details are available in table S2.

We then asked whether the percentage of A_dark_ spermatogonia is altered in the crypto samples. Assessment of the A_dark_ spermatogonia among all spermatogonia (MAGEA4^+^) indeed unveiled a significant reduction of this cell type in crypto samples (Fig. 6E). Specifically, we found a general reduction in A_dark_ cells among the PIWIL4^+^, UTF1^+^, FGFR3^+^, as well as GFRA1^+^ spermatogonia (Fig. 6, F to I). Finally, we questioned whether the A_dark_ reduction was dependent on the most advanced germ cell stage present in the respective tubules. Quantifications revealed that the number of A_dark_ spermatogonia in the crypto samples was reduced independently of the spermatogenic state of the tubules (Fig. 6J), suggesting that the loss of this cell type is a general process in crypto tissues.

### Altered EGR4 and HOXC9 expression in crypto State 0-0A spermatogonia

To assess potential changes in gene expression observed in the undifferentiated spermatogonia of crypto samples, we employed trajectory-based differential expression analysis (tradeSeq) *(18*). For this, cells were aligned along the latent time (fig. S6A) and then subdivided into six groups containing equal cell numbers (knotgroups) using seven knots (fig. S6B). We focused our attention on the most undifferentiated spermatogonia, State 0 and 0A, which are mostly represented by knotgroups 1 and 2.

Comparing knotgroup 1, which includes 78.2% of State 0 spermatogonia, between the normal and crypto datasets revealed 21 up- and 12 downregulated genes (Fig. 7A) involved in regulating cellular development and differentiation (Fig. 7B); among those genes, we found *EGR4* itself and 16 genes belonging to the EGR4 regulon (*ANXA11*, *CEP131*, *CRIP2*, *EGR4*, *IDH2*, *NUDT14*, *PDLIM4*, *PLPP2*, *PODXL2*, *PSIP1*, *RAC3*, *SH2B2*, *SMARCD3*, *SPINT1*, *ST3GAL4,* and importantly *UTF1*) (Fig. 7C and fig. S6D). We had already identified an enhanced activation of the EGR4 regulon in undifferentiated spermatogonia of the crypto samples (Fig. 4C), highlighting this regulon and gene as important regulators of spermatogonia. We then analyzed its expression in the spermatogonial states and found that, while expression of *EGR4* was restricted to State 0 in the normal situation, in the crypto dataset it showed a prolonged expression, being present also in States 0A and 0B (Fig. 7D). Afterwards, the expression of all EGR4-regulated genes was examined according to the spermatogonial states, confirming their modulation (fig. S6D). Interestingly, in the crypto group, we found a reduced expression of *UTF1* in State 0, 0A, and 0B spermatogonia (Fig. 7D).

**Fig. 7.**
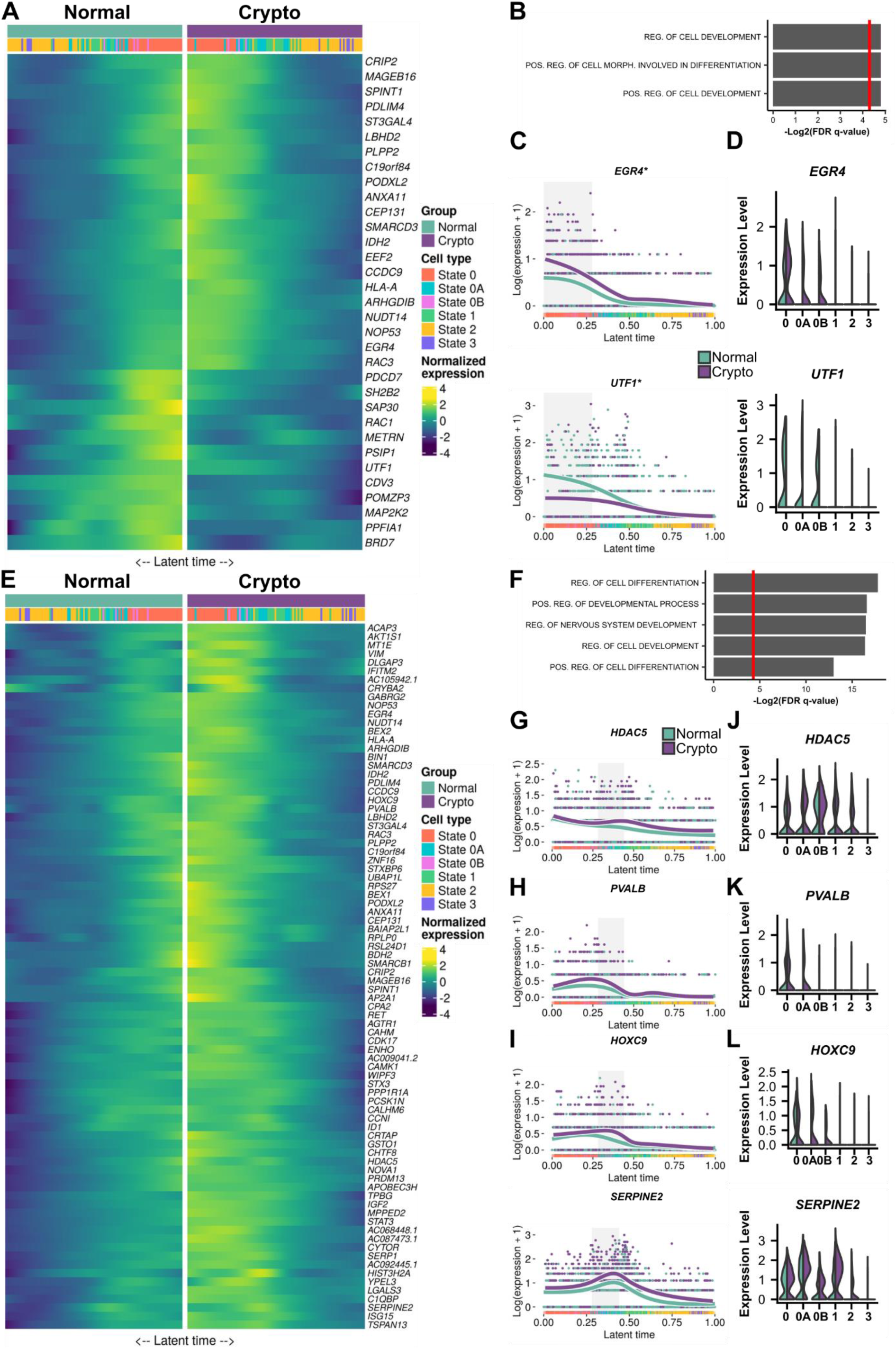
Crypto State0 and 0A spermatogonia show enhanced expression of EGR4 and HOXC9. (**A**) Double heatmap showing the normalized expression of the 33 genes identified by tradeSeq analysis as differentially expressed in the knotgroup 1 of the crypto dataset. The cells are plotted along the latent time with the normal cells on the left side (teal) and crypto cells on the right side (purple). (**B**) The bar plot shows GO analysis of the 33 differentially expressed genes in the knotgroup 1. The results showed an enrichment in GO terms related to cellular differentiation and development. The red line on the bar plot represents p=0.05. (**C**) Line plots showing the expression of *EGR4* and *UTF1* in the normal (teal) and crypto (purple) cells along the latent time. The gray area highlights the cells belonging to the knotgroup 1. The plots show enhanced expression of *EGR4* and reduced expression of *UTF1* in the crypto knotgroup 1. (**D**) Double violin plots comparing the expression level of *EGR4* and *UTF1* in the spermatogonial states in the normal (teal, left) and crypto (purple, right) datasets. (**E**) Double heatmap showing the normalized expression of the 82 genes identified by the tradeSeq analysis as differentially expressed in the knotgroup 2 of the crypto dataset. The cells are plotted along the latent time with the normal cells on the left side (teal) and crypto cells on the right side (purple). (**F**) The bar plot shows the GO analysis of the 82 differentially expressed genes in the knotgroup 2. The results showed an enrichment in GO terms related to cellular differentiation and development. The red line on the bar plot represents the p=0.05. (**G-I**) Line plots showing the expression along the latent time of *HDAC5* (G; regulated by EGR4 only), *PVALB* (H; regulated by HOXC9 only) and *SERPINE2* and *HOXC9* (I; regulated by EGR4 and HOXC9) in the normal (teal) and crypto (purple) cells. The gray area highlights the cells belonging to the knotgroup 2. The plots show enhanced expression of all the genes in the crypto knotgroup 2. (**J-L**) Double violin plots comparing the expression level of *HDAC5* (J; present in the EGR4 regulon only), *PVALB* (K; present in the HOXC9 regulon only), *SERPINE2* and *HOXC9* (L; present in both the EGR4 and HOXC9 regulons) in the normal (teal) and crypto (purple) cells along the latent time. The plots show enhanced expression of the all the genes in the crypto State 0, 0A, 0B spermatogonia.

For knotgroup 2, representing mostly State 0A cells, we identified 82 upregulated genes in the crypto cells (Fig. 7E), most of which are involved in cell differentiation (Fig. 7F). Again, *EGR4* was one of the differentially expressed genes, together with *HOXC9*, which we had already identified in the crypto samples as an activated regulon in undifferentiated spermatogonia (Fig. 4D). Among the differentially expressed genes in knotgroup 2, we found 25 genes that are regulated only by EGR4 (Fig. 7G, fig. S6C and fig. S7A), 1 gene that is regulated only by HOXC9 (Fig. 7H), and 5 genes that are regulated by both transcription factors, including *HOXC9* itself (Fig. 7I and fig. S7C). Among the 25 EGR4-regulated genes in knotgroup 2, 12 were also upregulated in knotgroup 1 (fig. S6C).

Assessing the expression of these genes according to the spermatogonial states, we observed that, similar to *EGR4*, *HOXC9* also displayed a prolonged expression in the crypto samples compared to the normal situation, where it is exclusively expressed in State 0 (Fig. 7L). This altered expression pattern was also observed for additional genes regulated by one or both of these transcription factors (Fig. 7, J to L and fig. S7, B to D), indicating that crypto samples show persistent expression of State 0 genes in undifferentiated spermatogonia (State 0 through 1).

## Discussion

In this work we pioneered the study of human male germ cell defects using state-of-the-art approaches. These led us to identify novel target genes and mechanisms for explaining the etiology of infertility. At cellular level we unveiled major alterations of the spermatogonial compartment with an increase in the number of the most undifferentiated (PIWIL4^+^) spermatogonia but a reduction in the morphologically defined A_dark_ reserve stem cells. Interestingly, these alterations are contingent on a prolonged expression of EGR4 in crypto samples. Also, we identified receptor-ligand interactions modulating the interplay between spermatogonia and their microenvironment.

ScRNA-seq analyses revealed six spermatogonial states based on transcriptional profile, in line with published data *(11*). Interestingly, we found a representation of the same states in both normal and crypto groups. Although the total number of spermatogonia remained unchanged, the states were represented in different proportions in the two groups. The most prominent finding was an increase, in the crypto group, of PIWIL4^+^ State 0 cells, which have been suggested as the origin of the spermatogonial differentiation process *(9, 11, 19)*. The increase of PIWIL4^+^ cells occurred concomitantly with a reduction of the UTF1^+^ cells, which we observed both at transcriptional and protein level. Moreover, we identified changes in transcriptional dynamics of crypto spermatogonia, as visualized by cells in State 0, uniformly pointing towards the direction of 0A and 0B in the RNA velocity analysis. As, PIWIL4^+^ and UTF1^+^ spermatogonia are mostly quiescent *(9, 11, 20, 21)*, we hypothesize that these altered cell proportions are a consequence of a change in transcriptional profiles of spermatogonial subpopulations and not due to changes in cell proliferation (Fig. 8A). We speculate that this increase in PIWIL4^+^ spermatogonia is the consequence of the reduced spermatogenic efficiency in cryptozoospermic men, however we cannot exclude that the failure to progress to more differentiated states might be the cause.

**Fig. 8.**
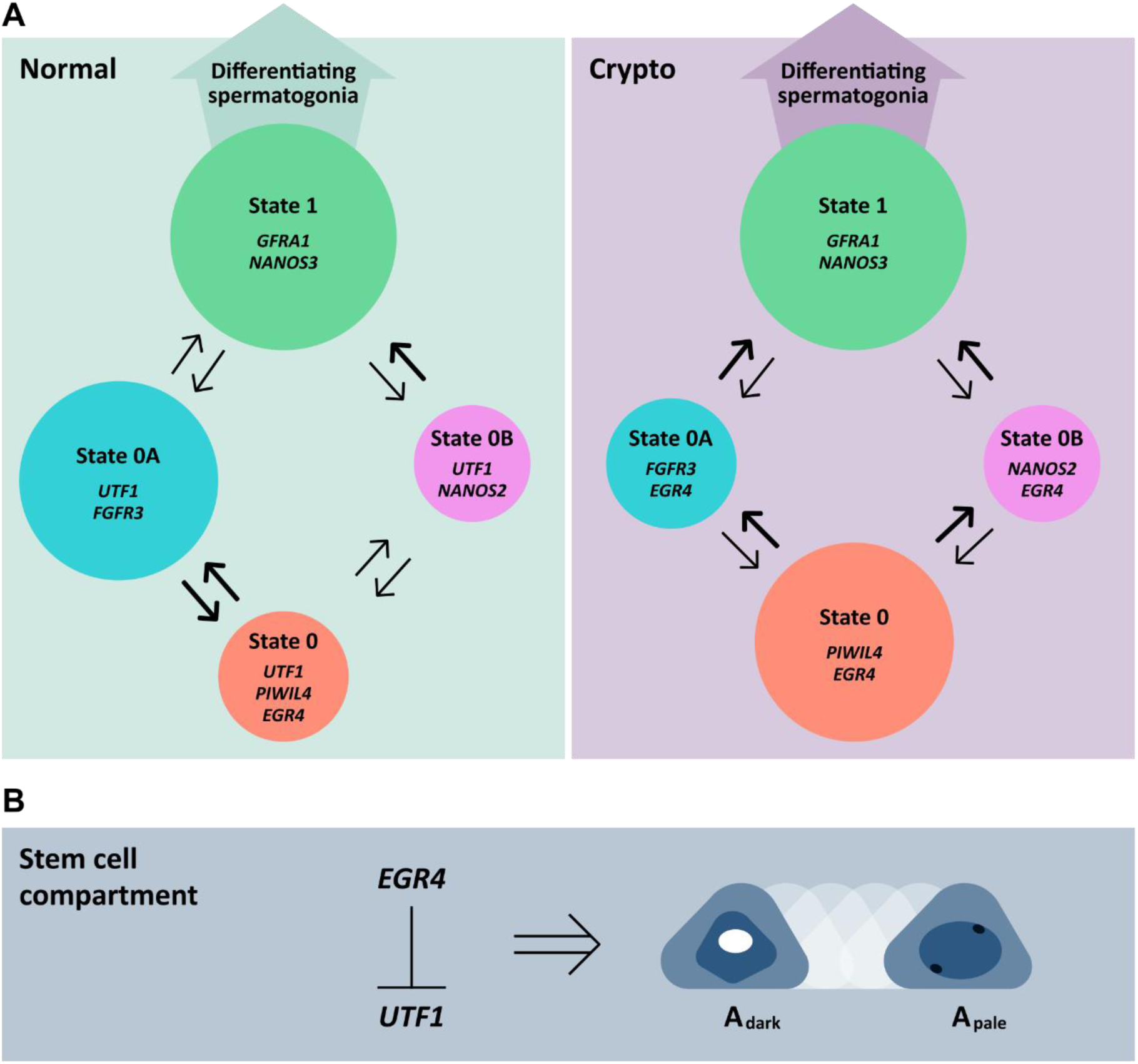
Model of the human spermatogonial compartment in normal and crypto samples. **(A)** The circles represent the State 0 (Red), State 0A (Blue), State 0B (Magenta) and State 1 (Green) spermatogonia. The thickness of the black arrows indicates the proportion of cells changing their expression profile. (**B)** Suggested mechanism: the EGR4 mediated UTF1 downregulation induces the chromatin remodeling resulting in the transition from A_dark_ to A_pale_ morphology.

We identified the transcription factor EGR4 as one of the gatekeepers regulating the change in transcriptional profiles of spermatogonia in the crypto group. Supporting this hypothesis we found EGR4 exclusively expressed in spermatogonia at protein level and in the most undifferentiated spermatogonial state (State 0) at transcriptional level, in accordance with recent studies *(9, 11)*. Importantly, trajectory-based differential expression analysis showed prolonged expression of EGR4 in spermatogonial States 0, 0A, and 0B in crypto samples. Moreover, we found enhanced activation of the EGR4, ASCL2, HOXC9 and DLX5 regulons in undifferentiated spermatogonia of crypto tissues. These four genes, including *EGR4* itself, contain a putative binding site for EGR4 in their promoter region suggesting EGR4 as an upstream regulator. The crucial role of EGR4 in the germline is corroborated by data demonstrating that it controls multiple differentiation steps during mouse spermatogenesis *(22*). Furthermore, *EGR4* gene sequence variations as well as altered gene expression profiles have been identified in men with impaired spermatogenesis *(23, 24*). Importantly, EGR4, ASCL2, HOXC9 and DLX5 have also been assigned pivotal roles in other cell systems. Interestingly, in small cell lung cancer EGR4 regulates proliferation through induction of DLX5 expression among other genes *(25*). ASCL2 activation is involved in restoring the intestinal stem cell system following damage *(26*) while HOXC9 and DLX5 are involved in neuronal and osteogenic cell differentiation processes, respectively *(27, 28*). These genes are highly relevant for cell proliferation and differentiation processes in multiple stem cell systems and therefore dysfunction in these pathways may present a link between the germline and other cell systems crucial for general health.

One of the genes presenting a putative binding site for EGR4 is *UTF1*, whose expression negatively correlates with that of *EGR4*. UTF1 is a chromatin-associated protein expressed in human undifferentiated, quiescent spermatogonia *(20, 21*). It is involved in embryonal stem (ES) cell differentiation *(29*) and acts as transcriptional repressor *(30*). Knock-down of UTF1 in ES cells results in extensive chromatin decondensation *(31*). Remarkably, we found a general reduction of the A_dark_ spermatogonia - characterized by highly condensed chromatin *(6, 7*) - in the crypto group. Marker profiling of A_dark_ spermatogonia shows that almost 80% are UTF1^+^. We therefore hypothesize that the reduction in UTF1 expression leads to a change in chromatin condensation resulting in the transition from an A_dark_ to an A_pale_ morphology (Fig. 8B). Thereby at cellular level, these cells with a more open chromatin are maintained in a ‘ready to react’ status. The price of this change in operating status is the reduction of the reserve stem cells (A_dark_), which under normal conditions ensure the recovery of sperm production following a gonadotoxic insult, by their ability to transform into mitotically active A_pale_ spermatogonia *(32*). This alteration of the stem cell pool may therefore place crypto patients at a disadvantage in the case of gonadotoxic insult. Moreover, the over-recruitment of the reserve spermatogonia into mitotically active spermatogonia may not be sustainable in the long term and is likely to lead to further quantitative and qualitative deterioration of germ cells.

The A_dark_ morphology is shared by spermatogonial subpopulations characterized by the expression of PIWIL4, FGFR3, and GFRA1, which define distinct states at transcriptional level (Fig 8A). Intriguingly, only 5.8% of the PIWIL4^+^ cells showed A_dark_ morphology. These findings indicate that multiple spermatogonial states are able to display A_dark_ spermatogonial morphology, in line with previous data *(33*), and also that A_dark_ spermatogonia may not be the most undifferentiated spermatogonia in the human testis. This presents a paradigm shift in the field.

The fact that the stem cell potential of the A_dark_ spermatogonia will remain untested until it is possible to purify these cells for use in transplantation assays in primates (including the human) constitutes a limitation of this study. Unfortunately, due to profound species differences, including the lack of A_dark_ reserve stem cells in rodents, these cannot be used as a model for mechanistic studies.

The microenvironment is pivotal for SSC behavior *(34, 35*). We identified multiple changes to the SSC microenvironment in the crypto group. Importantly, PMCs, perivascular cells, macrophages, and immune cells showed increased numbers and enhanced expression of chemokines. These changes suggest the presence of a proinflammatory microenvironment in crypto testes. In addition to the changes in the somatic compartment, we identified a reduced number of spermatocytes from the pachytene stage onwards in crypto samples. Intriguingly, this was accompanied by an enhanced interaction between FGF2 - produced by pachytene and diplotene spermatocytes - and FGFR1 and 3 - present on spermatogonia. These interactions were solely significant in the crypto group. The effect of FGF2 on human spermatogonia remains largely unknown, however a pro-survival role for FGF2 has been shown *(36*). Gain of function mutations in FGFR2 and 3 have been found in spermatogonia supporting the survival of these cells and inducing their clonal expansion *(37*). We suggest that FGF2-FGFR3 pathway may be relevant for maintaining the size of the human spermatogonial compartment in the crypto situation.

The revelation of regulators of the human spermatogonial stem cell compartment and its properties provides enticing possibilities. We recommend screening the genes highlighted in this manuscript for pathogenic variants. This may help to decipher genetic defects affecting spermatogonia, and thus help to resolve additional cases of male infertility. Finally, this will result in improved counselling of infertile men about the chances for sperm retrieval prior to surgery. In addition, it will help avoid unnecessary surgical interventions, identify potential health risks for their offspring, and establish preventive measures regarding their own general health risks. Lastly, due to the potential long term exhaustion of the stem cell pool crypto patients may prospectively benefit from cryopreservation of sperm prior to active family planning.

## Materials and Methods

### Human testicular biopsies

Adult human testis samples (n=34) were obtained from patients undergoing surgery for microdissection testicular sperm extraction (mTESE) or histological evaluation at the Department of Clinical and Surgical Andrology (University Hospital in Münster, Germany). One additional testicular sample was excised from each patient for this study after written informed consent. Ethical approval was obtained for this study (Ethics Committee of the Medical Faculty of Münster and State Medical Board no. 2008-090-f-S).

### Patient cohort selection

We subjected six testicular tissues (normal=3; crypto=3) to scRNA-seq. We selected patient samples with qualitatively and quantitatively normal spermatogenesis (normal), and with cryptozoospermia (crypto). Clinical workup prior to surgery included full physical evaluation, hormonal analysis (including luteinizing hormone (LH), follicle stimulating hormone (FSH) and testosterone (T)) *(38*), semen analysis *(39*), and genetic analyses (including karyotype and screening for azoospermia factor (AZF) deletions). Exclusion criteria were known genetic causes of infertility, acute infections, testicular tumors, and a history of cryptorchidism. Normal patients were selected from those diagnosed with obstructive azoospermia. These patients had no sperm in their ejaculate, normal testicular volume, and normal FSH levels. Age-matched cryptozoospermic patients had a sperm concentration <0.1 million/ml (cryptozoospermia) in the ejaculate, a reduced testicular size, and, in most cases, elevated FSH levels. All patients had spermatozoa in their TESE samples, regardless of histological results. Detailed clinical information about all the patients is available in table S1. Moreover, we added independent samples for proteome and histological analyses to further expand the sample cohort. Due to sample limitation, only 2 of the 3 crypto samples could be included in the proteome analysis.

### Exome sequencing

The exomes of the three cryptozoospermic men were sequenced. Briefly, target enrichment was performed by SureSelect QXT Target Enrichment Kit according to the manufacturer’s protocol using the capture libraries Agilent SureSelect Human All Exon Kit V6. Sequencing was performed on the Illumina NextSeq®500 system. The data was evaluated for rare (minor allele frequency [MAF] in gnomAD <0.01), likely pathogenic variants (stop-, frameshift- and splice site-variants) in 170 candidate genes previously reported to be associated with impaired spermatogenesis according to Oud et al. *(40*). (for gene list and detected variants see table S9). Additionally, the recently published genes ADAD2, M1AP, MSH4, RAD21L1, RNF212, SHOC1, STAG3, SYCP2, associated with non-obstructive azoospermia, were screened. Only variants with a coverage >10 and detected in genes that quantitatively impair spermatogenesis were considered for further analyses. Variants in recessive genes were only evaluated if at least two were identified in the same individual. For putative dominant and X-linked genes, variants that were also found in fertile controls were not considered for further analyses.

### Preparation of single cell suspensions

Single cell suspensions were obtained using a two-step enzymatic digestion as previously described *(41, 42*). A total of 20 000 cells were used for ploidy analysis, 12 000 cells were prepared for scRNA-seq and 50 000 cells were stored at −80°C for proteomic analysis.

### Ploidy analysis

Ploidy analysis was performed on 20 000 cells of each sample prior to scRNA-seq. After centrifugation, the cells were incubated in the dark for 30 minutes with a solution containing 50 µg/ml Propidium Iodide (Sigma-Aldrich, Cat# P4170), 1 mg/ml bovine serum albumin (Sigma-Aldrich, Cat# A9647), 0.1% Triton X-100 (Sigma-Aldrich, Cat# 93443), and 10 µg/ml RNase A (Sigma-Aldrich, Cat# R6513) in phosphate-buffered saline (PBS). After incubation, approximately 10 000 cells were analyzed for each sample using a Beckman Coulter CytExpert QC Flow Cytometer. Debris was defined based on forward and side scatter and excluded from the analysis. The cells analyzed for DNA content were measured at 617 nm and assigned to the following categories according to staining intensity: 1) haploid cells (1C: spermatids), 2) diploid cells (2C: spermatogonia, somatic cells (e.g. Leydig peritubular and Sertoli cells)), 3) “double diploid” cells (4C: spermatocytes).

### ScRNA-seq library preparation and sequencing

12 000 cells per sample (normal=3; crypto=3) were suspended in MEMα at a concentration of 500 cells/µl and loaded onto the Chromium Single Cell A Chip. Library preparation was performed following the kit instructions (Chromium Single Cell Kit [v2 chemistry]). Briefly ∼6 000 cells per sample were captured by the 10x Genomics Chromium controller, after cDNA synthesis, 12-14 cycles were used for library amplification. The resulting libraries were quantified using the Lab901 TapeStation system (Agilent) before shallow sequencing (NextSeq-550 sequencer, Illumina) for quality control. The final, deep sequencing was performed on a NovaSeq6 000 sequencer (Illumina), using 2 × 150 bp paired-end sequencing.

### ScRNA-seq and proteome analyses

The scRNA-seq and proteome analyses are described in detail Supplementary Materials and Methods.

### PAS, immunohistochemical and immunofluorescence staining

Testicular biopsies were fixed in Bouin’s solution overnight and then washed in 70% ethanol. The tissues were paraffin-embedded and sectioned at 5 µm. Tissue sections were dewaxed with AppiClear (Applichem, Cat# A4632.2500), rehydrated through decreasing ethanol concentrations, and washed in distilled water. For periodic acid-Schiff/hematoxylin (PAS) staining, the slides were first incubated in 1% PA (Sigma-Aldrich, Cat# 1.005.240.100) and then in Schiff’s reagent (Sigma-Aldrich, Cat# 1.090.330.500). Cell nuclei were counterstained with Mayer’s hematoxylin solution (Sigma-Aldrich, Cat# 1.092.490.500). After rinsing in tap water, the slides were dehydrated through increasing ethanol concentrations, incubated in AppiClear solution, and closed with Merckoglas (Sigma-Aldrich, Cat# 1.039.730.001).

Immunohistochemical staining was performed as previously described *(42*). Briefly, after rehydration, the sections underwent heat-induced antigen retrieval using sodium citrate buffer pH 6.0. The endogenous peroxidase activity and the unspecific antibody binding were blocked using hydrogen peroxide (Hedinger, Cat# GH06201) and a solution containing goat serum (Sigma-Aldrich, Cat# G6767-100ML) and BSA, respectively. The sections were incubated overnight at 4°C with the primary antibodies diluted in the blocking solution. The following day, sections were incubated with a biotin-labeled secondary antibody and then with streptavidin-horseradish peroxidase. The peroxidase activity was detected using 3,30-diaminobenzidine tetrahydrochloride solution (Applichem, Cat# A0596.0001). The reaction was stopped with distilled water and nuclei were counterstained with Mayer’s hematoxylin. Finally, the slides were rehydrated with increasing ethanol concentrations, washed with AppiClear, and closed with Merckoglas. The sections were scanned with Precipoint M8 Microscope and Scanner (Precipoint, Freising, Germany) *(43*).

The immunofluorescent staining was performed as previously described *(21*). After rehydration, the sections underwent heat-induced antigen retrieval using sodium citrate buffer pH 6.0. After cooling to room temperature the tissues were permeabilized with Triton X-100, incubated with 1M glycine (Sigma-Aldrich, Cat# G7126-500G) and then covered with a blocking solution of tween (Sigma-Aldrich, Cat# 655205), BSA, and donkey serum (Sigma-Aldrich, Cat# S30-100 ml). The incubation of the primary antibody was performed overnight at 4°C in blocking solution. The following day the sections were washed and incubated for 1 hour with species specific secondary antibodies diluted in blocking solution. Slides were finally mounted with Vectashield Mounting Media with 4,6-diamidino-2-phenylindole as nuclear counterstain (Vector Laboratories, Cat# H-1200). Stainings were digitalized employing the Olympus BX61VS microscope and scanner software VS-ASW-S6 (Olympus, Hamburg, Germany). All antibodies are listed in table S10.

### Evaluation of the testicular histology

To score the spermatogenic status of each patient two PAS-stained sections from two independent biopsies per testis were evaluated using the Bergmann and Kliesch method *(44*). A score from 0 to 10 was assigned to each patient according to the percentage of tubules containing elongated spermatids. In the same sections, the percentage of seminiferous tubules containing spermatocytes or spermatogonia as the most advanced germ cell type was assessed, as well as Sertoli cell only and hyalinized tubules (tubular shadows).

### A_dark_, A_pale_, B spermatogonia identification

The A_dark_, A_pale_ and B spermatogonia were defined according to morphological criteria as previously published *(6, 7*). Briefly, the A_dark_ spermatogonia presented a spherical or slightly ovoid nucleus containing uniformly dark stained chromatin with an unstained central rarefaction zone. A_pale_ spermatogonia had an ovoid nucleus containing very weakly stained chromatin and 1-3 deeply stained nucleoli attached to the nuclear membrane. Finally, B spermatogonia showed a spherical nucleus with granulated and weakly stained chromatin and intensely stained nucleoli detached from the nuclear membrane.

### Histological quantifications and statistics

Quantification of stained cells following immunofluorescence analyses was performed with Fiji/ImageJ *(45*). The number of ACTA2^+^ and MUSTN1^+^ blood vessels was evaluated in two independent sections of normal (n=12) and crypto (n=13) patients and normalized per area (mm^2^). To discern between the ACTA2^+^ seminiferous tubules and the ACTA2^+^ blood vessels for each stained section a consecutive section was stained with the endothelial marker VWF. A blood vessel was defined as such provided that it was positive for VWF in the consecutive section.

Each immunohistochemical quantification was performed on normal (n=6) and crypto (n=6) samples using the Precipoint Viewpoint software (Precipoint, Freising, Germany). The number of MAGEA4^+^, PIWIL4^+^, FGFR3^+^, GFRA1^+^, NANOS3^+^ and UTF1^+^ spermatogonia per tubule was assessed per round tubule. Tubules were considered round when the ratio between the two diameters was in the range of 1 - 1.5. The percentage of PIWIL4^+^, FGFR3^+^, GFRA1^+^, NANOS3^+^ and UTF1^+^ A_dark_ spermatogonia was evaluated counting 100 A_dark_ spermatogonia per sample and determining the proportion of each marker. In case of the crypto samples it was not always possible to reach 100 A_dark_ spermatogonia. To evaluate how many MAGEA4^+^, PIWIL4^+^, FGFR3^+^, GFRA1^+^ and UTF1^+^ spermatogonia were A_dark_, we counted 100 spermatogonia positive for each marker in each sample and then assessed the proportion of A_dark_ spermatogonia. Tubules of the crypto samples were subdivided into three categories according to the most advanced germ cell type present: spermatogonia, spermatocytes and spermatids. All the quantification results were plotted as box plots (center line: median; box limits: upper and lower quartiles; whiskers: 1.5× interquartile range; points: outliers). Normality and homoscedasticity tests were performed for all variables and differences between groups were assessed by parametric (t-test) or non-parametric tests (Mann-Whitney U test) as appropriate. Kruskal-Wallis rank sum test was used to compare three or more independent groups, followed by multiple pairwise comparisons. Statistical analysis and graphs were executed using R 4.0.0 and packages stats (v4.0.0) and ggplot2 (v3.3.1). Details regarding the number of samples or cells evaluated and the statistical analysis are provided in table S2.

## Supplementary material

Supplementary Materials and Methods

Fig. S1. Clustering analysis of normal and crypto datasets.

Fig. S2. DGE and GO analysis between the different germ cell clusters of the normal and crypto datasets.

Fig. S3. DGE and GO analysis between the different somatic cell clusters of the normal and crypto datasets.

Fig. S4. Differential regulon activation in the normal and crypto datasets.

Fig. S5. Cluster analysis and assignment of normal and crypto spermatogonial datasets.

Fig. S6. EGR4-regulated genes in the knotgroup 1 spermatogonia.

Fig. S7. EGR4- and HOXC9-regulated genes in knotgroup 2 spermatogonia.

Table S1. Clinical parameters.

Table S2. Histomorphometric and statistical analysis.

Table S3. Single cell RNA sequencing stats.

Table S4. Differential expression analysis results.

Table S5. Genes uniquely differentially expressed in germ or somatic cell clusters and relative gene ontology analysis.

Table S6. Differential gene expression analysis of perivascular cells compared toPMCs and of immune cells compared to all other cell clusters.

Table S7. List of the 403 regulons obtained from the Scenic analysis.

Table S8. List of regulons obtained from the Scenic analysis plotted in the heatmap in Fig. 4.

Table S9. Genes with at least one potentially pathogenic variant described according to Oud et al. 2019.

Table S10. Reagents & Software.

## Acknowledgements

We thank Klaus Redmann for excellent technical support in PI analyses of testicular samples. We thank Nicole Terwort for excellent technical support in processing of testicular tissues, and we thank Heidi Kersebom and Elke Kößer for histological evaluation of testicular tissues. We also thank Sabine Forsthoff for excellent support in endocrinological measurements and Willi Kramer, Fotozentrale University Hospital of Münster, for the illustration in Fig. 1A.

## Author Contributions

S.D.P., T.T., S.L. and N.N. designed the study, performed data analysis, data interpretation and wrote the manuscript; H.C.A.D. performed low-input proteome analyses; M.W., F.T. performed whole exome sequencing and data analyses; X.L., G.M.Z.H. provided the infrastructure and expertise for single cell RNA sequencing and prepared sequencing libraries; T.T., M.D. performed bioinformatic analyses of single cell RNA sequencing datasets, L.M.S.K., S.D.P. performed experiments; J.F.C., S.K. supervised clinical patient care and documentation in the database, performed surgeries and were responsible for clinical and laboratory testing; J.W., S.S. provided expertise for histomorphometrical analyses and data interpretation; N.N. coordinated the project. All authors approved the final version of the manuscript.

## Data and materials availability

All mass spectrometry proteomics data were deposited to the ProteomeXchange Consortium (http://proteomecentral.proteomexchange.org) *(46)* via the PRIDE partner repository. The data will be provided upon request.

The sequencing data generated during this study as well as Alevin counts, integrated counts and a cell metadata table are available at the NIH Gene Expression Omnibus (GEO) under accession number GSE153947 (www.ncbi.nlm.nih.gov/geo/query/acc.cgi?acc=GSE153947). The data will be provided upon request.

## Supplementary Material

### Material and Methods

#### ScRNA quantification and UMI/CB correction

Alevin *(47*) was used to deduplicate and quantify the single-cell sequencing data and to discern the valid cellular barcodes (CB) from the background noise. The Gencode *(48*) human reference v30 was used as transcriptomic reference, together with the flags --useCorrelation and -- chromium in the quantification. Additionally, the --expectCells parameter was set to 6 000 (50% of the initial loading of 12 000 cells) to help the CB knee finding method. This approach resulted in a mean of 4 783 cells (SD: 587) per sample and a mean/median number of 2 460 (SD: 799) expressed genes per cell. All software is listed in table S10.

#### Normalization, sample integration, dimensional reduction and label transfer

Alevin counts were processed with Seurat *(14, 49*). To reduce initial noise, genes expressed in less than 0.1% of the cells were excluded, together with cells containing fewer than 200 counts. In total, only 9 cells were removed by this filter. The analysis of mitochondrial content in regard to the total measured expression revealed 471 cells (1.6%) with a strong mitochondrial effect (>25% of mitochondrial counts), 306 of them (65%) later identified as Sertoli cells. It is unknown at the moment if this high mitochondrial content is a sign of cell death or a normal feature of Sertoli cells, therefore these cells were not excluded from the analysis. Mitochondrial content and total RNA counts were used as a regression variable in the normalization procedure sctransform *(50*), with the number of variable features set to 5 000. Anchor-based sample integration was performed on the normalized counts, setting the number of features in the anchor finding process to 5 000. We integrated normal and crypto samples (n=3, each) in three ways, resulting in an integrated dataset of all normal samples, an integrated dataset of all crypto samples and an integrated dataset consisting of all six samples. As proposed by the Seurat developers, scaling was disabled in the FindIntegrationAnchors() method, due to the prior normalization by sctransform. All available genes were kept in the integrated dataset, irrespective of their usage as anchors or not. Integrated counts were scaled and centered for principal component (PC) analysis and dimensional reduction. For each scaled dataset, 100 PCs were calculated using the most variable genes. Elbow-Plots of the PCs were generated to determine the inflection point, confirming 25 PCs for further analysis. A shared nearest neighbor graph was constructed based on the PCs. Cell clusters were identified via the smart local moving algorithm *(51*) with a resolution parameter of 1 100 random starts and a maximum of 100 iterations. For data visualization, the non-linear dimensional reduction method UMAP *(52*) was used based on the specified 25 PCs.

To enable an unbiased verification of the cluster identities, the top marker genes per cluster were computationally determined with the FindAllMarkers() function, testing with MAST *(53*) only for positive marker genes with a minimal expressed-in fraction of 30% (p < 0.001). These computational markers were used with known cell type markers from the literature *(9–12)* to carefully assign cluster identities. Following the marker-based cluster identity assignment in the normal dataset, identity labels were transferred to the crypto dataset utilizing the Seurat label transfer functionality. Imputed cell identity labels were carefully inspected and again confirmed by marker expression. The same clustering and dimensional reduction approach was performed on the subgroup of spermatogonial cells of the normal and crypto dataset.

#### Differential gene expression analysis

Intra- and inter-dataset differential gene expression analysis was performed using MAST *(53*). The original Alevin expression counts (after default Seurat normalization) were used to avoid bias by the scaling effect of sample-wise sctransform normalization or sample integration. Filter criteria for differentially expressed genes (DEG) were: default log fold change threshold of 0.25, FDR-corrected p-value below 0.01 and expression in at least 10% of the cells of one comparison group.

#### PAGA trajectory inference

Partition based graph abstraction (PAGA) *(54*) from the Scanpy package *(55*) was used for trajectory inference. The neighborhood graph was based on the top 25 PCs from Seurat with the number of nearest neighbors set to 20. Cell identity labels were used as partition categorical for the PAGA construction. Based on the PAGA, a new UMAP embedding was computed to better reflect the global topology, using 100 optimization iterations (epochs).

#### RNA velocity estimation and embedding

RNA velocity is the time derivative of the measured mRNA abundance (mature spliced/ nascent unspliced transcripts) and allows to estimate the future developmental directionality of each cell. As the abundance estimation relies on genomically aligned nascent reads, it is not possible to use the transcriptomic pseudo-alignment of Alevin. Therefore, we used the 10x Genomics Cell Ranger pipeline to create a genomic BAM file for each sample. Subsequently, aligned reads were used separately as input for velocyto *(17*), as well as the 10x Genomics genomic annotation file and UCSC expressed repeats annotation file. Velocyto analysis of each sample was limited to cell barcodes validated by Alevin. Due to differences in the barcode correction methods of Alevin and Cell Ranger, RNA velocity could not be estimated for a total of 24 cells (0.08% of total cells). Per sample abundance estimates were combined with the loompy package, analogous to the scheme in the Seurat integration step. Combined abundance data was merged with the PAGA anndata object using scVelo *(56*). Abundance estimates were normalized and filtered, requiring a minimal abundance of 5 per gene, simultaneously in the spliced and unspliced counts. First and second order moments were calculated utilizing the first 30 PCs and 30 neighboring cells from the neighbor graph. The likelihood-based dynamical model from scVelo was applied, recovering full splicing kinetics before estimating the velocities. In addition to the velocity graph and the projection of the velocities into the UMAP embedding, a gene-shared latent time was calculated. The latent time is solely based on the transcriptional dynamics and represents the internal clock of a cell. For the latent time calculation, a cell from the most undifferentiated cell type was used as a root cell, if applicable.

#### Gene regulatory network inference

The python version of the SCENIC *(16*) pipeline was used for gene regulatory network inference. Briefly, SCENIC links cis-regulatory sequences to single-cell gene expression data to predict interactions between transcription factors (TFs) and target genes. Pairs of genes and co-expressed TFs were identified with the GRNBoost2 algorithm from Arboreto *(57*), using the standard normalized Alevin counts. Additionally, a list of 1390 curated human TFs was used, provided in the SCENIC repository. Co-expression modules derived from this analysis were pruned to remove indirect targets and false positives (using human motifs v9 and hg38 databases from cisTargetDB). In accordance with the SCENIC publication we refer to modules with significant motif enrichment of the correct upstream regulator as ‘regulons’. Cells with enriched expression for genes in a regulon were marked as active for this specific regulon. Additionally, a Jensen-Shannon divergence based regulon specificity score *(58*) (RSS) was used for ranking purposes.

As we experienced highly variable results for different SCENIC runs on the same data, we choose a strategy to increase the stability of the adjacency calculation, i.e. the target gene and transcription factor co-expression measurement and weighting. Because of its nondeterministic nature, we decided to perform this calculation 10 times and only carry over gene–TF links that appeared in each of the runs. This “meta-adjacency” table, consisting of the median weight per link, was used in subsequent SCENIC steps.

#### Differential expression analysis along latent time

We utilized the R package tradeSeq *(18*) to perform differential expression analysis along the common gene-shared latent time between the spermatogonial cells of the normal and crypto datasets. Standard normalized Alevin counts were filtered, excluding genes with a total expression below 10 (excluding 28% of 33 679 genes). For each gene, a negative binomial generalized additive model was fitted to seven knots along the latent time, in accordance to the results of evaluateK(). The knots are equally distributed among the cell density along the trajectory, with the first and last knot representing the minimal and maximal trajectory value, respectively. Knots were comprised to knot groups, where all cells between adjacent knots are included. We focused on identifying genes differentially expressed exclusively in a single knot group. For this, we adopted the stageR *(59*) two-staged testing scheme, performing a whole-trajectory patternTest() for the screening stage and a earlyDETest() for each knot group in the confirmation stage. All tests were performed against a foldchange of log_2_ (1.2) with an overall false discovery rate of 0.05. The fitted distributions of the significant genes of each earlyDETest() were clustered, using 100 cluster points and a minimal cluster size of 20, except for knotgroup 1, where a minimal cluster size of 10 was used.

#### Correlation analysis of spermatogonial cell types

Cell types of the subset of spermatogonial cells were assigned based on the conjunction of markers of spermatogonial states, as described by Guo et al. and Sohni et al. *(9, 11*). To asses that the cell type assignment worked as expected, correlation analysis of spermatogonial states between our dataset and the datasets of Guo and Sohni was performed. Spearman correlation of common genes was based on an “average” cell, consisting of the genewise mean of all cells of the specific state.

#### Cell-cell communication analysis

Analysis of cell-cell communication was performed with CellphoneDB *(15*) (Version 2.1.2), using the “statistical_analysis” method. Ligands and receptors were required to be expressed in at least 50% of the cells (--threshold) and the number of iterations was set to 10 000. Significant interactions (p < 0.05) were subsequently visualized.

#### Gene ontology and pathway analysis

Gene ontology analysis was performed using the Gene Set Enrichment Analysis (GSEA) (https://www.gsea-msigdb.org/gsea/index.jsp). Pathway analysis was performed using Ingenuity Pathway Analysis (Qiagen).

#### Sample preparation and LC-MS/MS analysis

Single cell suspensions obtained by digestion of a biopsy (see “Single cell suspension preparation” section) from either normal (n=5) or crypto patients (n=4) were prepared for bottom-up mass spectrometry analysis, combining a one-pot sample preparation method (iST-NHS; Preomics) with isobaric tandem mass tags (TMT) labeling and subsequent stepwise reversed-phase fractionation at high pH (Thermo Fisher Scientific, Pierce, Cat# 84868). Cell lysis, digestion and labeling were performed according to the manufacturer’s protocol, except for the two samples containing 1.0×10^6^ cells (assuming a protein content of ∼200 µg), where volumes of buffers and reagents were adjusted accordingly (N1, N2). In addition, to support complete degradation of released chromatin, 1 µl of Benzonase was added to each sample after the initial lysis/denaturation step and incubated for 10 minutes at room temperature (RT). TMT labeling was performed following digestion by directly adding the TMT labeling reagent resuspended in 41 µl dry acetonitrile to the corresponding digest (RT, 500 rpm, 1hr). Prior to quenching the labeling reaction with 5% hydroxylamine, a “label check” was performed by analyzing 1 µl of each sample by mass spectrometry. A second mass spectrometry analysis was performed by mixing 1 µl of each sample to obtain normalization factors for the correct 1:1 ratio across all channels. Following desalting and lyophilization of the mixture, labeled peptides were resuspended in 300 µl of 0.1% TFA and submitted to a reversed-phase fractionation procedure at high pH on spin columns according to the manufacturer’s instructions (5% - 50% acetonitrile, 0.1% triethylamine). This resulted in 8 fractions (plus desalted flowthrough and 2 wash fractions) that were dried down in an Eppendorf concentrator before being resuspended in 0.1% formic acid for mass spectrometry analysis. All samples were measured twice as technical replicates. Samples were measured on an Easy nLC 1200 system coupled to a Q Exactive HF MS via a Nanospray Flex ion source (ThermoFisher Scientific). Peptides were dissolved in buffer A (0.1% formic acid) and separated on a 25 cm column, in-house packed with 1.9 µm C18 beads (Reprosil -Pur C18 AQ, Dr. Maisch) using a multi-linear gradient from 5%-18% buffer B (80% acetonitrile; 0.1% formic acid) and from 18%-40% B in 55 min each, followed by an increase to 60% B in 10 min, a final washout for 7 min at 90% B and re-equilibration at starting conditions (100% buffer A; flow rate 250 nl/min). The Q-Exactive HF mass spectrometer was operated in data-dependent acquisition mode (spray voltage 2.1 kV; column temperature maintained at 45°C using a PRSO-V1 column oven (Sonation, Biberach)). MS1 scan resolution was set to 120 000 at m/z 200 and the mass range to m/z 350-1 600. AGC target value was 3E6 with a maximum fill time of 50 ms. Fragmentation of peptides was achieved by higher-energy collisional dissociation (HCD) using a top15 method (MS2 scan resolution 60 000 at 200 m/z; AGC Target value 1E^5^; maximum fill time 108 ms; isolation width 1.3 m/z; normalized collision energy 32). Dynamic exclusion of previously identified peptides was allowed and set to 30 s, singly charged peptides and peptides assigned with a charge > 8 were excluded from the analysis. Data were recorded using Xcalibur software (ThermoFisher Scientific).

#### MS data analysis and quantification

Raw MS files were processed using MaxQuant (version 1.6.6.0) *(60*). Identification of peptides and proteins was enabled by the built-in Andromeda search engine by querying the concatenated forward and reverse mouse Uniprot database (UP000005640_9606.fasta; version from 04/2019) including common lab contaminants. Allowed initial mass deviations were set to 7 ppm and 20 ppm, respectively, in the search for precursor and fragment ions. Trypsin with full enzyme specificity and only peptides with a minimum length of 7 amino acids was selected. A maximum of two missed cleavages was allowed; the ‘match between runs’ option was turned on. Carbamidomethylation (Cys) was set as fixed modification, while Oxidation (Met) and N - acetylation at the protein N-terminus were defined as variable modifications. For peptide and protein identifications, a minimum false discovery rate (FDR) of 1% was required. First, the list of identified proteins was filtered and potential contaminants, reverse hits derived from the target-decoy search as well as proteins that were identified only by a single modified peptide were removed. Only proteins identified by at least one unique peptide were retained for further analysis. Next, a sample loading normalization step was performed using an R script (https://pwilmart.github.io/TMT_analysis_examples/CarbonSources_MQ.html), which was adapted to our own experimental setup. The list of proteins containing the normalized reporter ion intensities was imported into the Perseus bioinformatics tool. Proteins that did not show reporter ion intensities in all 10 TMT channels were removed, and reporter ion intensities were log_2_ transformed. To screen for differentially expressed proteins, a two-sided Student’s t-test was performed using permutation-based FDR calculation as truncation (FDR = 0.05), also taking into account the contribution of the difference of the means (s0 = 0.1).*(61*)

## Supplementary figures

**Fig. S1.**
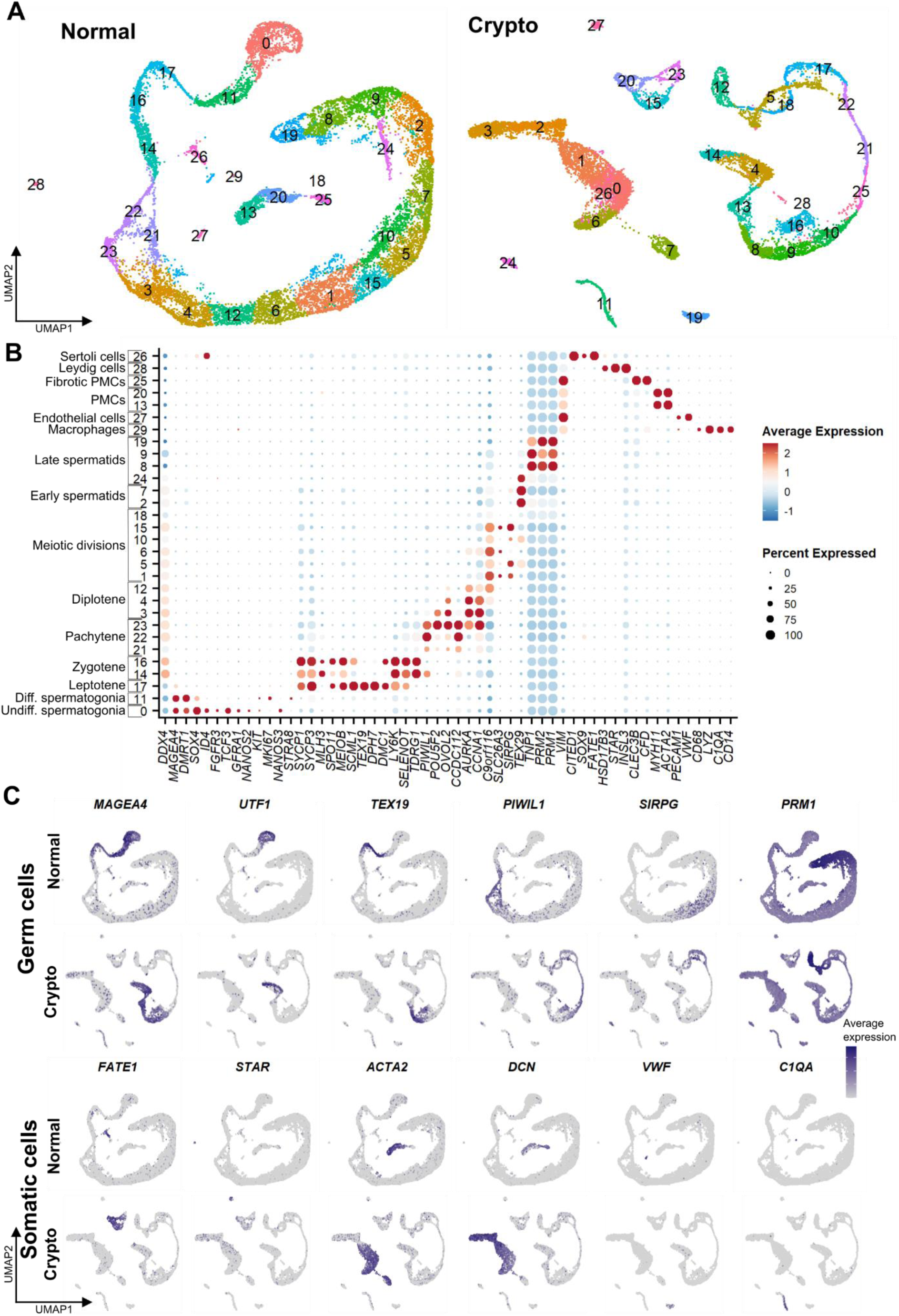
Clustering analysis of normal and crypto datasets. (**A**) Uniform manifold approximation and projection (UMAP) plot of the clustering analysis of the integrated (left) normal (15 546 cells, 30 clusters) and (right) crypto (13 144 cells, 29 clusters) datasets. (**B**) Dot plot showing the relative expression of 55 marker genes in the 30 normal clusters. Cell identity was assigned according to marker expression. The color and size of the dots represent the average expression and the percentage of cells expressing each marker in a cluster, as indicated in the key. (**C**) Feature plots showing the expression of germ cell and somatic cell marker genes in the normal and crypto datasets.

**Fig. S2.**
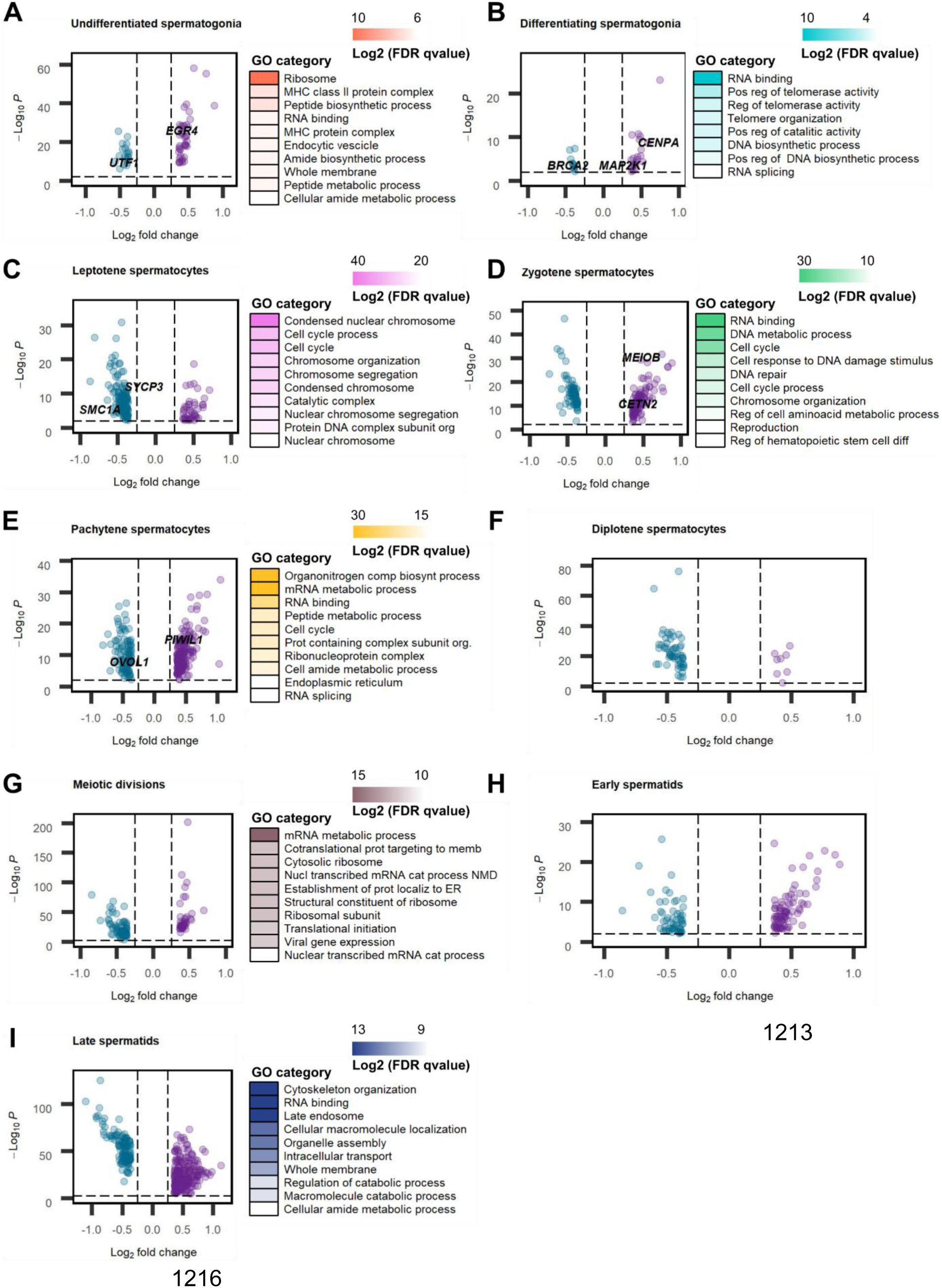
DGE and GO analysis between the different germ cell clusters of the normal and crypto datasets. (**A**) Volcano plot (Left) and gene ontology (GO) analysis (Right) of the 61 genes uniquely differentially expressed in the undifferentiated spermatogonial cluster. (**B**) Volcano plot (Left) and GO analysis (Right) of the 32 genes uniquely differentially expressed in the differentiating spermatogonial cluster. (**C**) Volcano plot (Left) and GO analysis (Right) of the 164 genes uniquely differentially expressed in the leptotene spermatocyte cluster. (**D**) Volcano plot (Left) and GO analysis (Right) of the 180 genes uniquely differentially expressed in the zygotene spermatocyte cluster. (**E**) Volcano plot (Left) and GO analysis (Right) of the 271 genes uniquely differentially expressed in the pachytene spermatocyte cluster. (**F**) Volcano plot of the 66 genes uniquely differentially expressed in the diplotene spermatocyte cluster. (**G**) Volcano plot (Left) and GO analysis (Right) of the 121 genes uniquely differentially expressed in the meiotic division cluster. (**H**) Volcano plot of the 142 genes uniquely differentially expressed in the early spermatid cluster. (**I**) Volcano plot (Left) and GO analysis (Right) of the 405 genes uniquely differentially expressed in the late spermatid cluster. See table S5 for the statistical details.

**Fig. S3.**
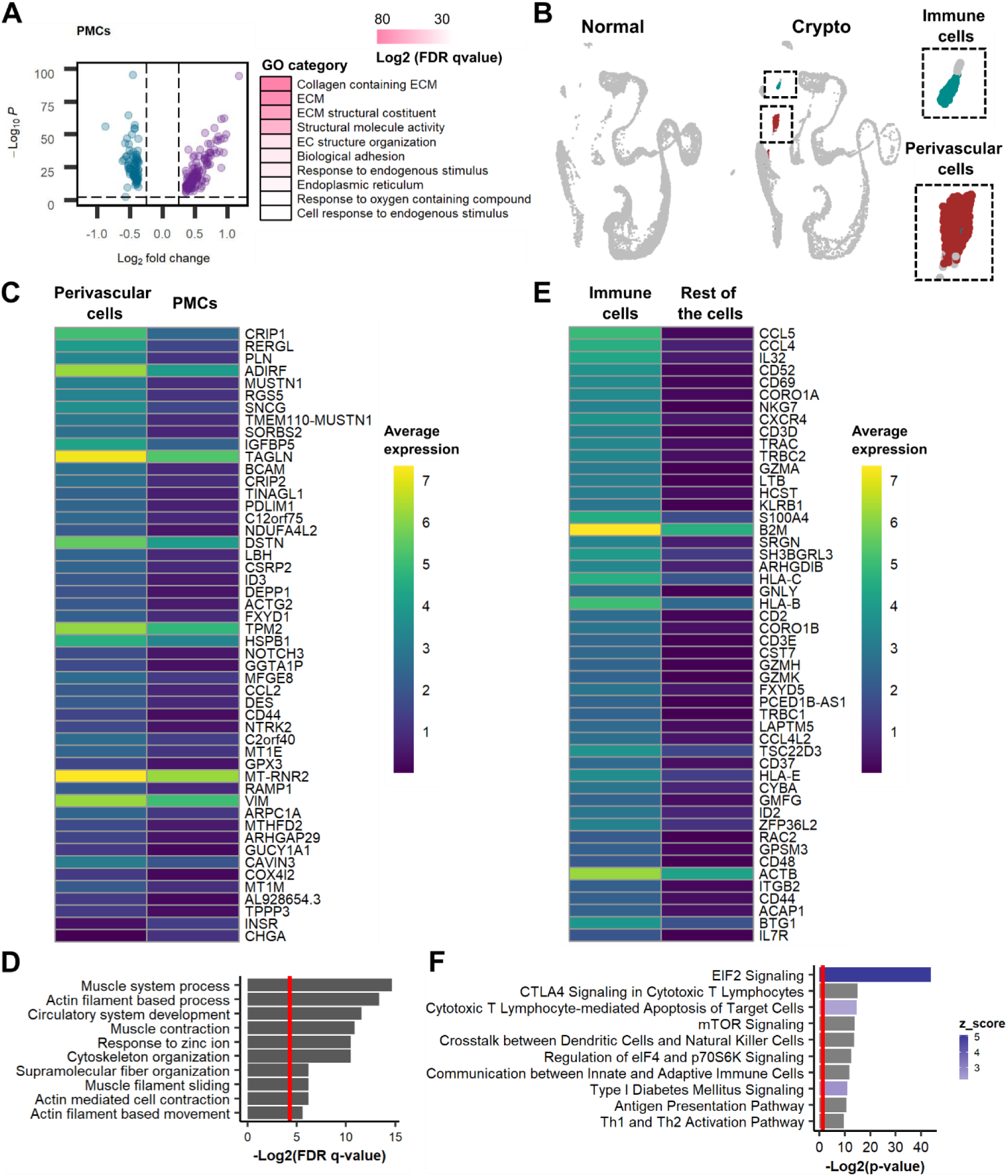
DGE and GO analysis between the different somatic cell clusters of the normal and crypto datasets. **(A)** Volcano plot (Left) and gene ontology (GO) analysis (Right) of the 232 genes uniquely differentially expressed in the PMC cluster. See Supplementary information, Table S5 for the statistical details. **(B)** Uniform manifold approximation and projection (UMAP) plot of the normal (Left) and crypto (Right) datasets showing the unique presence of the peritubular (Red) and immune (Green) cell clusters in the crypto dataset. **(C)** Heatmap showing the differential gene expression analysis between the perivascular cell clusters and the PMCs in the crypto dataset. The analysis identified 48 differentially expressed genes (DEG), including MUSTN1, with a higher expression in perivascular cells (table S6). **(D)** The bar plot shows the GO analysis of the 48 DEG found in perivascular cells. The results were enriched in GO terms related to muscle and cytoskeletal contraction. The red line on the bar plot represent p=0.05. **(E)** Heatmap showing the first 50 genes of the differential gene expression analysis between immune cells and the rest of the cells in the crypto dataset. A total of 195 genes showed higher expression in cluster 24 (A complete list of the DEG is available table S6). **(F)** The bar plot shows the pathway analysis of the 195 DEG. The results were enriched in immune cell associated pathways. The red line on the bar plot represents the p=0.05. The z_score represents the activation level of each pathway.

**Fig. S4.**
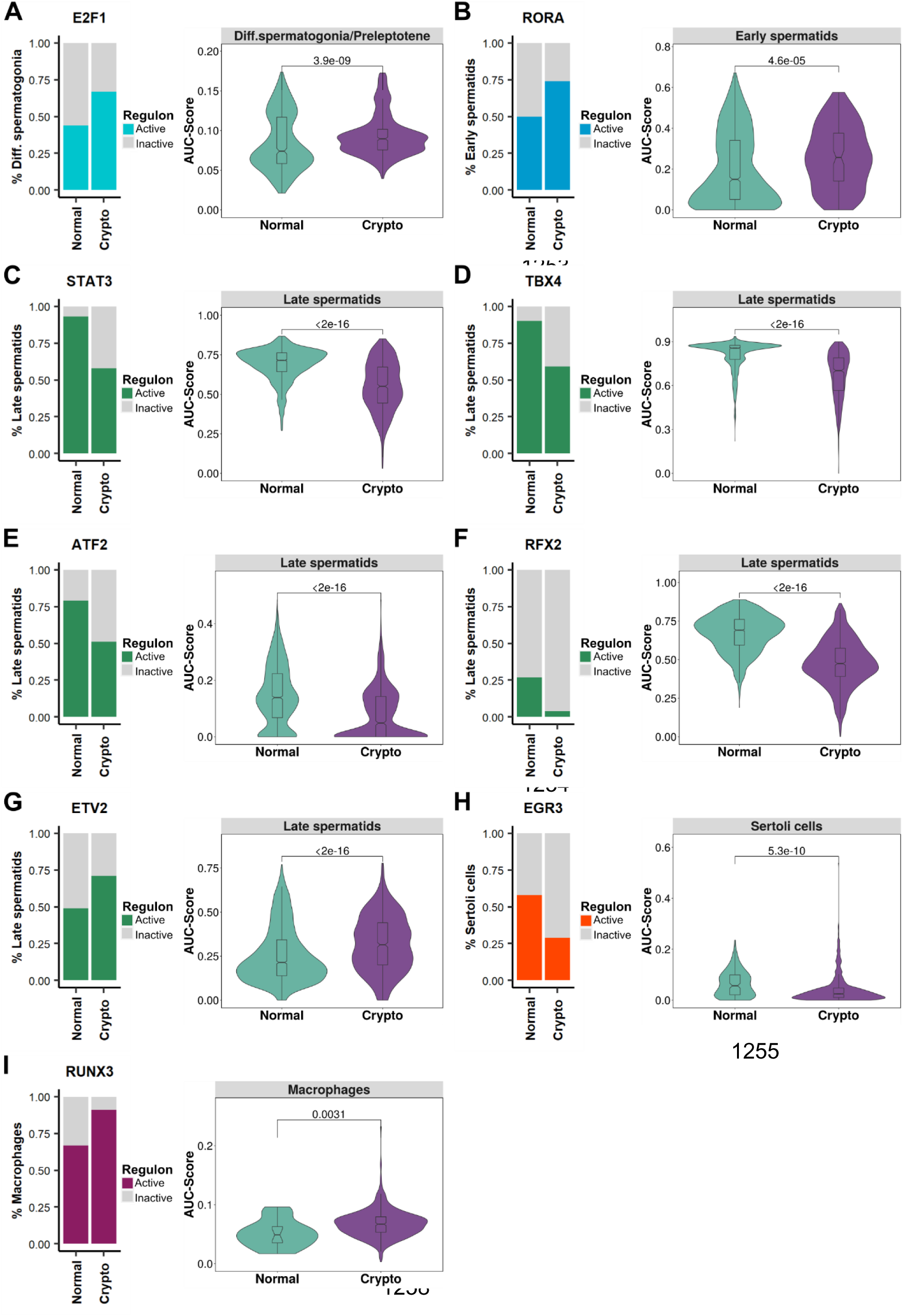
Differential regulon activation in the normal and crypto datasets. (**A-I**) Left panel: Stacked bar plot comparing the proportion of cells with active regulons. Right panel: Violin plots comparing the AUC score of the regulons in the normal and crypto dataset. The regulons are organized as follow: (**A**) E2F1 in the differentiating spermatogonia/preleptotene, (**B**) RORA in the early spermatids, (**C**) STAT3, (**D**) TBX4, (**E**) ATF2, (**F**) RFX2 and (**G**) ETV2 in the late spermatids, (**H**) EGR3 in the Sertoli cells and (**I**) RUNX3 in the macrophages. A significant change was found for all the nine regulons in the crypto dataset.

**Fig. S5.**
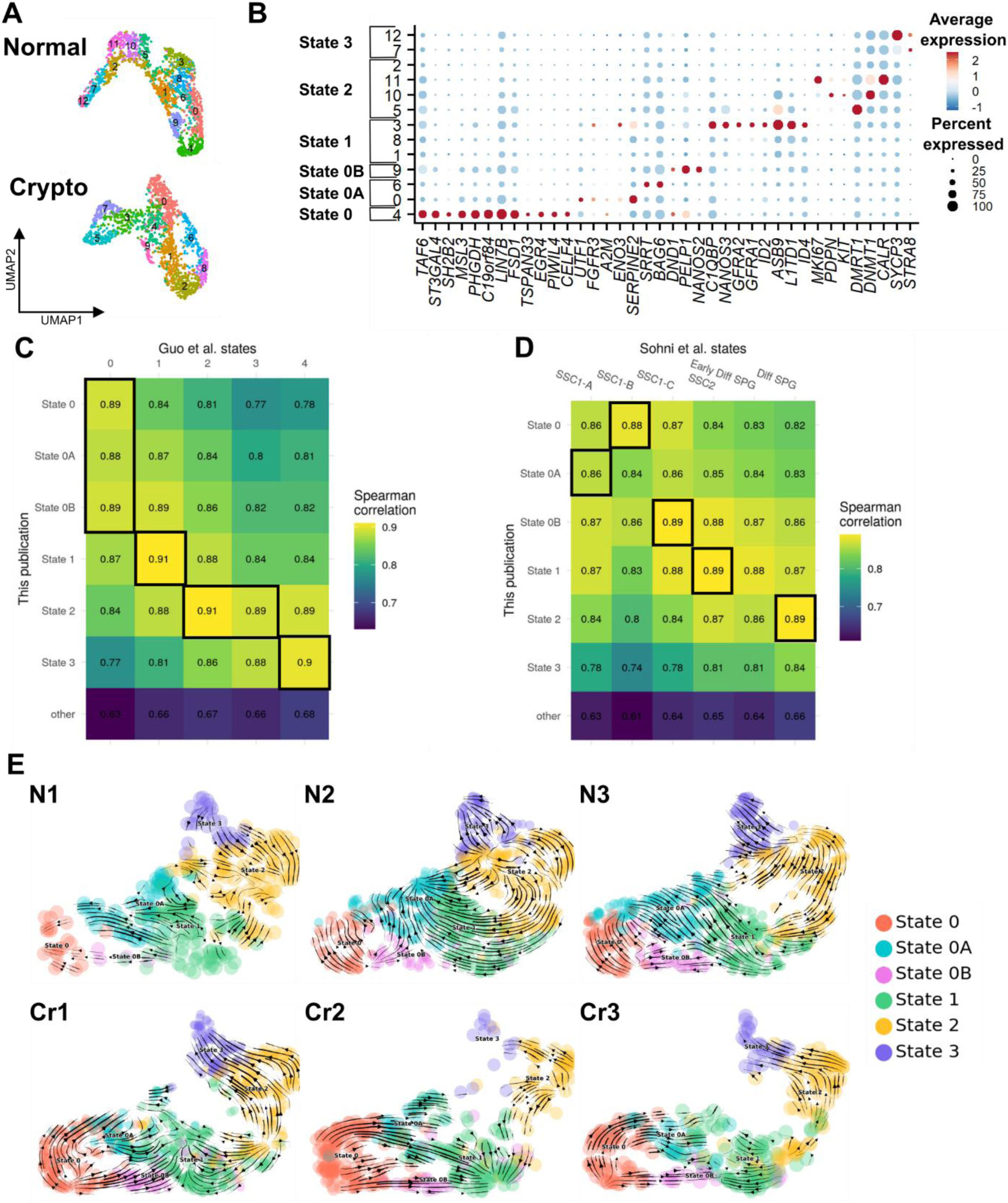
Cluster analysis and assignment of normal and crypto spermatogonial datasets. **(A)** Uniform manifold approximation and projection (UMAP) plot of the integrated normal (upper panel, 13 clusters) and crypto (lower panel, 10 clusters) spermatogonial datasets. The cells are color coded according to the cluster. **(B)** Dot plot showing the relative expression of selected marker genes in the 13 normal clusters. The color and size of the dots represent the average expression and the percentage of cells expressing each marker in a cluster, as indicated in the key. According to the marker expression, six clusters were defined: State 0, 0A, 0B, 1, 2, and 3. **(C-D)** Correlation analysis comparing the transcriptome of the different spermatogonial states defined in this publication with those defined in Guo et al., 2018 **(C)** and Sohni et al., 2019 **(D)**. The ‘other’ state is comprised of all non-spermatogonial cell types available in this publication. The black boxes show the states where we expected the highest correlation in each state. **(E)** Contribution of each normal and crypto sample to the RNA velocity derived from the scVelo dynamical model visualized as streamlines in UMAP plots.

**Fig. S6.**
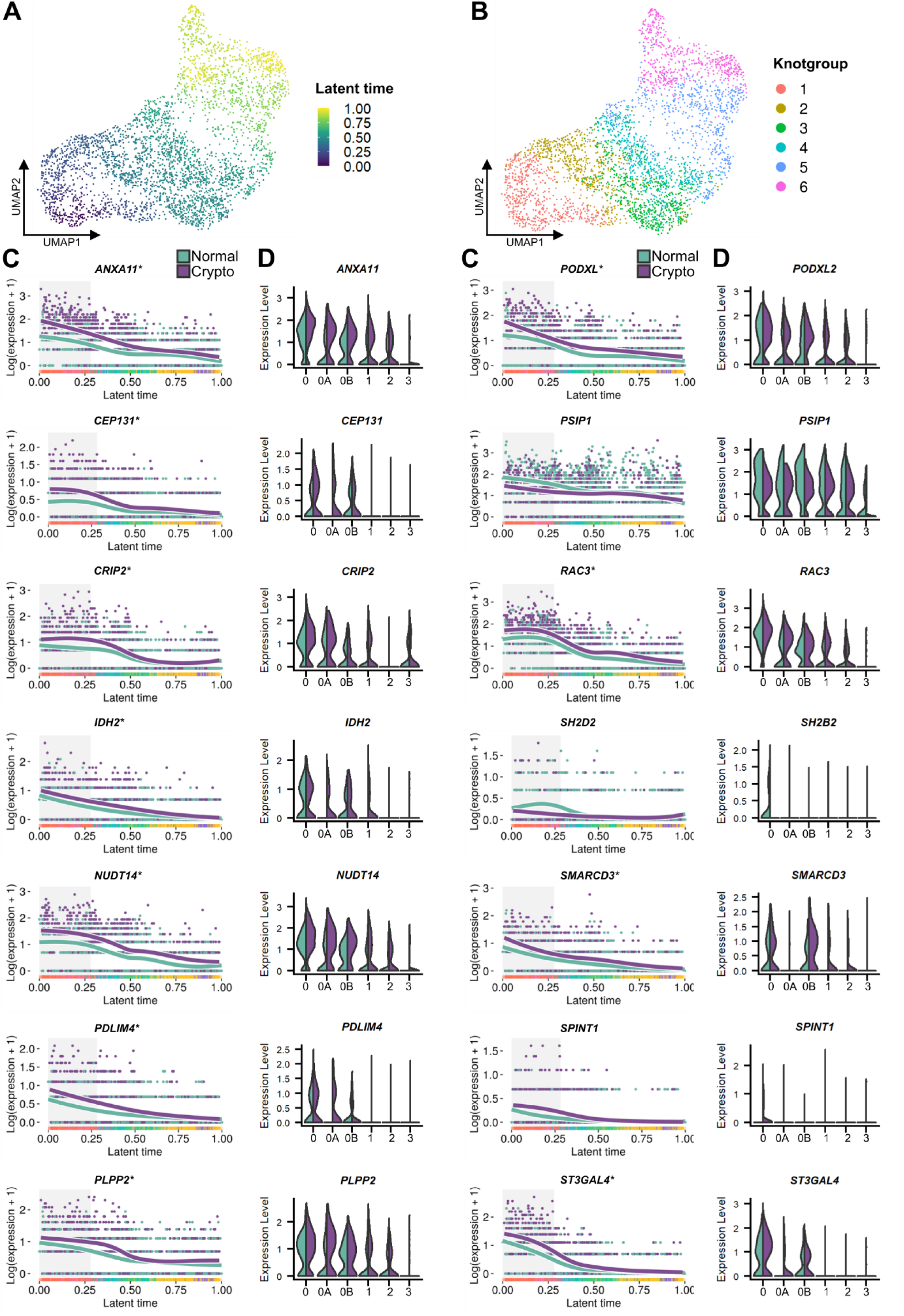
EGR4-regulated genes in the knotgroup 1 spermatogonia. (**A**) Uniform manifold approximation and projection (UMAP) plot showing the integrated normal-crypto spermatogonial dataset aligned along the latent time. The cells are color-coded according to their progression along the latent time. State 0 was set as starting point of the differentiation process. (**B**) UMAP plot showing the subdivision of the integrated normal-crypto spermatogonial dataset into six knotgroups. The cells are color coded according to their respective knotgroup. (**C**) Line plots show the expression along the latent time of the EGR4-regulated differentially expressed genes in normal (teal) and crypto (purple) spermatogonia included in the knotgroup 1: ANXA11, CEP131, CRIP2, IDH2, NUDT14, PDLIM4, PLPP2, PODXL2, PSIP1, RAC3, SH2B2, SMARCD3, SPINT1, and ST3GAL4. The gray area highlights the cells belonging to the knotgroup 1. The asterisk indicates EGR4-regulated genes that were also differentially expressed in knotgroup 2. (**D**) Double violin plots comparing the expression levels of EGR4-regulated ANXA11, CEP131, CRIP2, IDH2, NUDT14, PDLIM4, PLPP2, PODXL2, PSIP1, RAC3, SH2B2, SMARCD3, SPINT1, and ST3GAL4 in the spermatogonial states of the normal (teal) and crypto (purple) datasets.

**Fig. S7.**
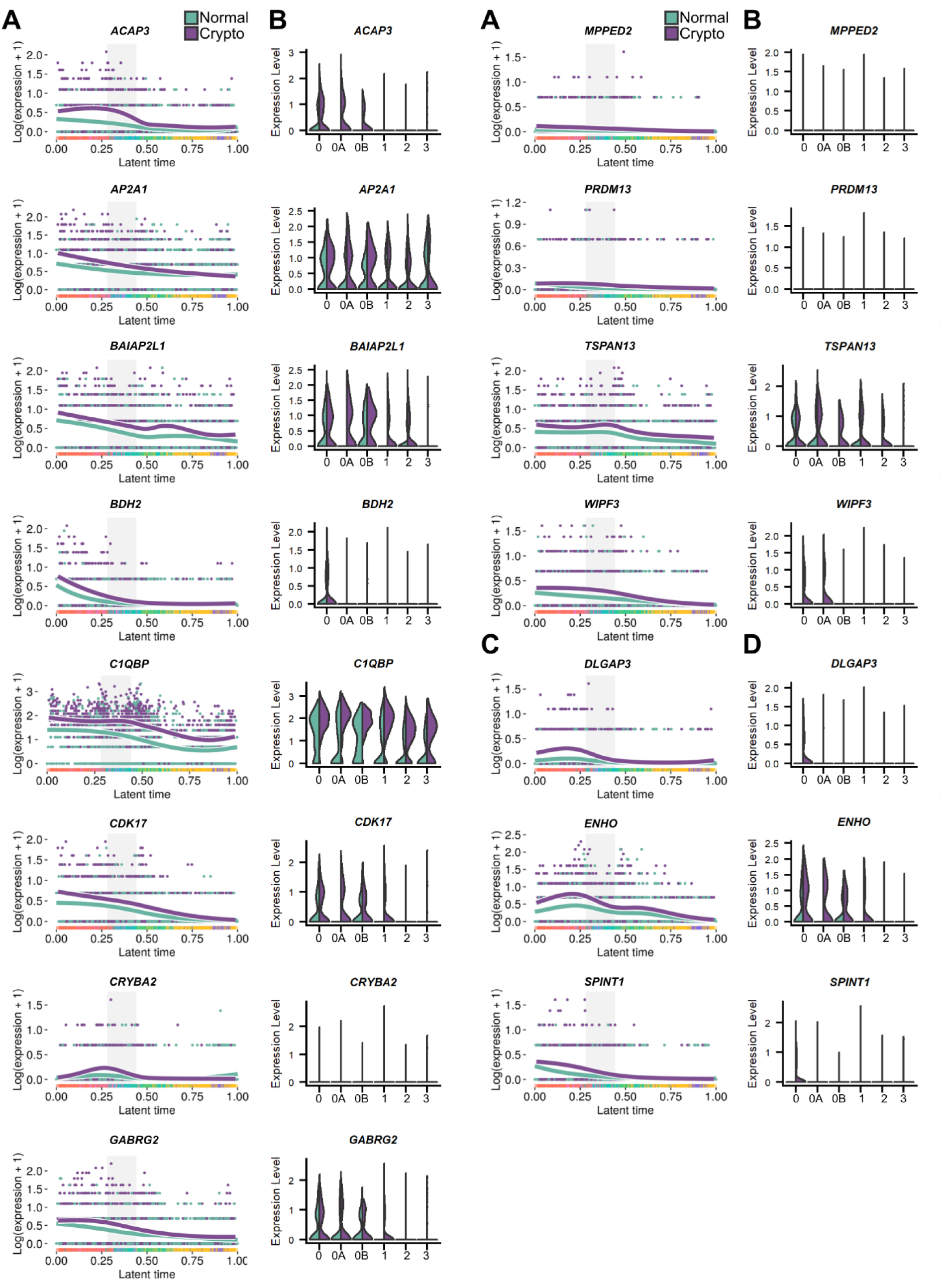
EGR4- and HOXC9-regulated genes in knotgroup 2 spermatogonia. (**A**) Line plots showing the expression along the latent time of EGR4-regulated, differentially expressed genes between the normal (teal) and crypto (purple) spermatogonia in the knotgroup 2: ACAP3, AP2A1, BAIAP2L1, BDH2, C1QBP, CDK17, CRYBA2, GABRG2, MPPED2, PRDM13, TSPAN13, and WIPF3. The gray area highlights the cells belonging to the knotgroup 2. (**B**) Double violin plots comparing the expression levels of EGR4-regulated genes ACAP3, AP2A1, BAIAP2L1, BDH2, C1QBP, CDK17, CRYBA2, GABRG2, MPPED2, PRDM13, TSPAN13, and WIPF3 between the normal (teal) and crypto (purple) spermatogonial states. (**C**) Line plots showing the expression along the latent time of the EGR4- and HOXC9-regulated, differentially expressed genes between the normal (teal) and crypto (purple) spermatogonia in knotgroup 2: DLGAP3, ENHO, and SPINT1. The gray area highlights the knotgroup 2. (**D**) double violin plots comparing the expression levels of EGR4- and HOXC9-regulated genes DLGAP3, ENHO, and SPINT1 between the normal (teal) and crypto (purple) spermatogonial states.

## Supplementary tables

Data files table S1-S10

